# Enhanced intercellular transfer of mitochondria from nuclear respiratory factor 1 (NRF1)-primed mesenchymal stem cells: towards creation of superior mitochondrial delivery hubs

**DOI:** 10.64898/2026.02.06.704090

**Authors:** Hyunho Lee, Pinar Atalay, Gherardo Baudo, Matteo Massaro, Zheng Yin, Elvin Blanco

**Affiliations:** Center for BioNanoengineering, Houston Methodist Research Institute, Houston, TX 77030; Department of Systems Medicine and Bioengineering, Houston Methodist Neal Cancer Center, Houston, TX 77030; Department of Medicine, Weill Cornell Medical College, New York, NY, 10065; Department of Cardiology, Houston Methodist DeBakey Heart and Vascular Center Houston Methodist Hospital, Houston, TX 77030

**Keywords:** Mitochondrial dysfunction, nuclear respiratory factor 1 (NRF1), mesenchymal stem cells (MSCs), mitochondrial biogenesis, mitochondrial transfer

## Abstract

Mitochondrial dysfunction is a pervasive hallmark of diverse diseases. In endothelial cells (ECs), oxidative stress, bioenergetic failure, and dysregulated mitochondrial dynamics (fusion-fission, mitophagy) damage the endothelium and promote vascular pathologies such as diabetes, atherosclerosis, and aging. Mitochondrial augmentation, via direct transplantation of isolated mitochondria or cell-to-cell transfer of the organelle, has emerged as a strategy to restore mitochondrial function in metabolically compromised cells. We recently established that overexpressing nuclear respiratory factor 1 (NRF1), a driver of mitochondrial biogenesis, in mesenchymal stem cells (MSCs) increases mitochondrial content and preserves mitochondrial function under senescence-inducing stress. Here, we advance NRF1-primed MSCs as enhanced mitochondrial hubs for intercellular mitochondrial delivery to cells undergoing mitochondrial dysfunction. We hypothesized that NRF1 overexpression engages mitochondrial transfer machinery, thereby enhancing both tunneling nanotube (TNT)- and extracellular vesicle (EV)-mediated mitochondrial transfer to stressed ECs, improving EC mitochondrial fitness and health. mRNA-mediated NRF1 priming of MSCs increased expression of proteins involved in mitochondrial motility and transfer, enhanced TNT formation, and increased production of mitochondria-containing EVs. Single-cell RNA sequencing (scRNA-seq) results show that NRF1 priming shifted MSCs into distinct transcriptional states, with NRF1-enriched clusters exhibiting coordinated upregulation of cell-adhesion/cytoskeletal connectivity programs and vesicle-fusion/trafficking pathways, features consistent with enhanced structural coupling and secretory transfer capacity. NRF1 priming increased TNT-like F-actin intercellular bridges in direct co-culture and elevated mitochondria-containing EV transfer in transwell assays, demonstrating augmented mitochondrial delivery through both contact-dependent and contact-independent routes. Consequently, recipient ECs displayed reduced mitochondrial ROS, preserved membrane potential, improved oxidative phosphorylation and ATP production, rebalanced mitochondrial dynamics of fusion-fission and mitophagy. NRF1-primed MSCs further attenuated oxidative stress-induced EC senescence and apoptosis. Together, these findings identify NRF1 activation as a mechanism to reprogram MSCs into high-capacity mitochondrial donors and support NRF1-driven mitochondrial hub engineering as a strategy to strengthen mitochondrial transfer-based therapies for diseases characterized by mitochondrial dysfunction.

## Introduction

Mitochondrial dysfunction is a central, convergent driver of various diseases characterized by oxidative stress. In endothelial cells (ECs), impaired oxidative phosphorylation (OXPHOS) and ATP depletion weaken energy-dependent barrier maintenance (e.g., cytoskeletal remodeling and tight-junction stability), while excess mitochondrial ROS (mtROS) amplifies inflammatory signaling and leukocyte-endothelium interactions that exacerbate vascular dysfunction [1–3]. In ocular vascular beds, including the choriocapillaris and deep retinal plexus, these bioenergetic and redox defects contribute to barrier breakdown, leukocyte adhesion, and maladaptive angiogenic responses implicated in age-related macular degeneration (AMD) and ischemic retinopathies [4, 5]. Similar defects (e.g., impaired nitric oxide signaling, junctional disassembly, and pro-inflammatory activation) also characterize microvascular disease in the brain and other organs [6]. Taken together, these observations motivate therapeutic strategies that restore mitochondrial function in ECs undergoing oxidative stress rather than solely suppress downstream inflammation.

Despite the recognized role of mitochondrial dysfunction in endothelial disease, therapies that reverse bioenergetic failure remain limited. Our previous work showed that mitochondrial augmentation, including direct mitochondrial transplantation into metabolically compromised cells, restores cellular energetics, reduces mtROS, and improves mitochondrial function [7–9], supporting a mitochondria-centric strategy across disease contexts. Mitochondrial augmentation can also occur endogenously via intercellular mitochondrial transfer in response to stress. For example, astrocytes donate functional mitochondria to neurons via tunneling nanotubes (TNTs), improving neuronal survival after injury [10]. TNTs are actin-based, transient intercellular conduits induced by oxidative or inflammatory stress that enable directed trafficking of whole mitochondria to metabolically compromised cells, restoring ATP production and limiting stress signaling [11, 12].

Beyond TNTs, accumulating evidence indicates that mitochondria can be trafficked via extracellular vesicles (EVs). EV-associated mitochondrial transfer is being recognized as a component of intercellular mitochondrial quality control and stress adaptation, enabling contact-independent, longer-range support of recipient cell bioenergetics [13–16]. Mesenchymal stem cells (MSCs) are well known for therapeutic effects mediated by immunomodulation and secretion of trophic factors [17], including a secretome (conditioned media, CM) that delivers growth factors, cytokines, lipids, and coding or non-coding RNAs (ncRNAs) that influence EC proliferation, migration, tube formation, and barrier integrity [18–20]. Importantly, MSCs have emerged as mitochondrial donor hubs capable of transferring functional mitochondria via TNTs and EVs to improve OXPHOS and ATP generation in injured ECs [14, 21, 22]. However, oxidative disease milieus can induce premature senescence and death in native MSCs, undermining their regenerative potential. ROS-driven stress impairs MSC expansion, promotes mitochondrial dysfunction, and accelerates senescent pathways [23–27]. Senescence can also reshape EV biology, altering EV yield/composition and, critically, mitochondrial-related cargo (including mitochondrial components and/or mtDNA-associated inflammatory signaling) [25, 28], which may diminish therapeutic consistency of MSC-EV products derived under stress.

These limitations motivate strategies that reinforce MSC mitochondrial fitness to improve durability and potency. Nuclear respiratory factor 1 (NRF1) is a transcriptional regulator of mitochondrial biogenesis that activates nuclear-encoded programs supporting mtDNA replication and transcription through its activation of mitochondrial transcription factor A (TFAM), increasing mitochondrial content [29, 30]. We previously established that NRF1 induction alleviates amyloid-β-associated mitochondrial toxicity by promoting mitochondrial augmentation [31], reinforcing NRF1 as a tractable node for reversing disease-relevant mitochondrial defects. We further showed that NRF1 induction in MSCs enhanced mitochondrial function and diminished senescence, establishing a route to generate stress-resilient donor cells with elevated mitochondrial biogenesis, an essential precedent for the present study [32].

Here, we extend this framework to the vascular interface and formalize the concept of MSC mitochondrial hubs – NRF1-primed MSCs with fortified mitochondrial networks and increased mitochondrial content to enhance endothelium repair through reinforced mitochondrial delivery. We hypothesized that NRF1 activation engages transfer machinery, thereby augmenting TNT- and EV-mediated mitochondrial transfer to oxidative stress-exposed ECs, restoring mitochondrial function and endothelial health. We show that NRF1 activation in MSCs increases mitochondrial biogenesis and content and is associated with higher expression of proteins implicated in mitochondrial motility and transfer, enhanced TNT formation, and increased production of mitochondria-containing EVs. Single-cell RNA sequencing (scRNA-seq) supports upregulation of gene programs linked to mitochondrial transfer. Functionally, increased mitochondrial transfer from MSCs to ECs was accompanied by reduced oxidative stress, improved cell respiration, and rebalanced mitochondrial dynamics. NRF1-primed MSCs attenuated oxidative stress-induced EC senescence and apoptosis, preserving EC function. By coupling intrinsic mitochondrial biogenesis to TNT- and EV-mediated intercellular mitochondrial transfer, this work defines a mechanistically grounded platform for endothelial remodeling and positions NRF1-primed MSC mitochondrial hubs as a strategy to restore vascular homeostasis in oxidative-stress vasculopathies.

## Materials and Methods

### Materials

Human NRF1 mRNA was purchased from TriLink Biotechnologies (San Diego, CA) Scrambled control (SCR) mRNA was provided by the Houston Methodist RNA Core (Houston, TX). Human bone marrow-derived mesenchymal stem cells (MSCs) were obtained from Lonza (Durham, NC). HUVECs were obtained from ATCC (Manassas, VA). Lipofectamine™ MessengerMAX™ reagent was purchased from Thermo Fisher Scientific (Waltham, MA). MSC basal medium and SingleQuotes^TM^ supplements were purchased from Lonza bioscience (Houston, TX). Human umbilical vein endothelial cells (HUVECs), vascular cell basal medium, and endothelial cell growth supplements were purchased from ATCC (Manassas, VA). Lentiviral constructs encoding mitochondria-targeted reporters rLV.EF1.AcGFP1-mito-9 (mito-AcGFP1) and rLV.EF1.mCherry-mito-9 (mito-mCherry) were obtained from Takara Bio (San Jose, CA). Protamine sulfate was purchased from Thermo Fisher Scientific, and puromycin was purchased from MilliporeSigma (Burlington, MA).

### Cell culture and maintenance

MSCs were cultured in MSC basal medium supplemented with MSCGM™ SingleQuots™. ECs were cultured in vascular cell basal medium supplemented with endothelial cell growth supplements. Cells were maintained at 37°C and 5% CO_2_ and used at passage 5 for all experiments. Cell numbers were determined at every passage, and cells were routinely seeded at 0.5 × 10^4^ cells/cm^2^ (MSCs) and 1 × 10^4^ cells/cm^2^ (ECs).

### Lentiviral labeling of mitochondria in MSCs and ECs

To primarily distinguish donor and recipient mitochondria during transfer assays, MSCs were transduced with mito-AcGFP1 and ECs were transduced with mito-mCherry. MSCs were seeded at 0.8 × 10^4^ cells/cm^2^ and ECs at 1.2 × 10^4^ cells/cm^2^. After 24 h, cells were transduced with lentiviral particles at MOI 10 in complete medium supplemented with protamine sulfate (100 µg/ml). After 24 h, medium was replaced with fresh complete medium and cells were cultured for an additional 48 h, followed by puromycin selection to generate stable mito-AcGFP1 MSCs and mito-mCherry ECs. mito-AcGFP1 MSCs were used to track mitochondria in EV-related experiments, and mito-AcGFP1 MSCs co-cultured with mito-mCherry ECs were used for intercellular mitochondrial transfer studies.

### MSC mRNA transfection

Prior to co-culture, MSCs were transfected with human NRF1 mRNA or SCR mRNA using Lipofectamine™ MessengerMAX™ according to our previously published protocol [32].

### Identification and quantification of TNTs

TNT identification and quantification were adapted from a published method [33]. Following F-actin staining, TNTs were imaged using a Nikon A1 confocal microscope (40x objective). TNTs were defined by all the following criteria: (i) suspended structures lacking substrate contact (confirmed by orthogonal views and z-stacks), (ii) thin membranous protrusions with diameter <1 µm, and (iii) continuous linear projections originating from one cell and extending uninterrupted to another cell, forming a direct intercellular bridge. Structures not meeting these criteria were excluded.

For quantitative analysis, TNTs and total cell numbers were manually counted using open-source ICY image analysis software (Institut Pasteur and France-BioImaging). Ten randomly selected fields per well were analyzed using identical imaging parameters across conditions. The number of TNTs was quantified by normalizing the TNT count to the total number of cells present in the corresponding 40x microscopic fields.

### Co-localization of mitochondria with EV markers in MSCs

To assess mitochondrial association with EV-related compartments, mito-AcGFP1 MSCs were seeded on glass coverslips, fixed with 4% paraformaldehyde (PFA), permeabilized, and immunostained with anti-CD63 (abcam, 1:200), followed by an Alexa Fluor 594-conjugated secondary antibody. Co-localization between AcGFP1 (mitochondria) and CD63 (EV/MVB; extracellular vesicle/multivesicular body marker) was quantified using the Coloc 2 plugin, and Manders’ colocalization coefficient was computed [34, 35].

### EV isolation

All reagents and buffers used for EV work were filtered through 0.1 µm pore-size filters (MilliporeSigma). EVs were isolated from conditioned media of non-transfected (NT), SCR- or NRF1-transfected MSCs. Six hours post-transfection, cells were washed twice with PBS to remove residual transfection reagents and serum. The medium was then replaced with MSC basal medium supplemented with 10% EV-depleted fetal bovine serum (FBS) to minimize bovine EV contamination. Supernatants were collected after 18 h and processed using a published protocol [36]. Conditioned media were centrifuged at 500 × g for 5 min to remove cells, then at 2,500 × g for 10 min to remove debris. The supernatant was then centrifuged at 18,000 × g for 30 min to pellet large EVs. EV pellets were washed once in filtered PBS and resuspended in PBS for downstream analyses.

### Confocal images of isolated EVs

EVs isolated from mito-AcGFP1 MSCs were incubated with Alexa Fluor 647-conjugated anti-CD63 antibody (Clone AD1, Cat# IC5048R, R&D Systems, 1:200) for 25 min at room temperature in the dark. PBS-only controls (no EVs) were processed in parallel to assess nonspecific/background signal. Samples were washed twice with filtered PBS, spotted onto glass slides, cover slipped, sealed, and imaged using a Nikon AX-R+NS PARC confocal system with a 100× oil immersion objective.

### Nanoparticle tracking analysis by ZetaView

EV concentration and size distribution were measured using a ZetaView Particle Metrix X30. EV samples were serially diluted in 0.1 µm-filtered PBS (1:10, then 1:100), for a final dilution of 1:1,000 prior to measurement. Samples were equilibrated to room temperature. Measurements were acquired in scatter mode using a predefined EV protocol with identical instrument settings across all samples. Particle concentration (particles/mL) and size distribution were computed in ZetaView software and averaged across multiple positions per sample.

### High-sensitivity spectral flow cytometric of EVs

To quantify mitochondria-containing EVs, isolated EVs were diluted in sterile filtered PBS and stained with Alexa Fluor 647-conjugated anti-CD63 antibody at 1:200 for 30 min in the dark. Data were acquired on a Cytek Aurora™ CS with the trigger set to violet side scatter (VSSC). Mitochondrial content was identified by intrinsic AcGFP1 fluorescence (blue-laser excitation), and CD63 by Alexa Fluor 647 (red-laser excitation). Samples were acquired at low flow to minimize coincidence/swarm detection and collected for a fixed duration (5 min) to enable absolute event comparison. Buffer-only controls and single-stain controls (EVs lacking mitochondrial label and/or without CD63 staining) were included to define background and spectral unmixing. Total EVs were defined as CD63⁺ events; mitochondria-containing EVs were defined as CD63⁺/AcGFP1⁺ dual-positive events. Data were analyzed in Floreada.io with identical gating across NT, SCR, and NRF1 groups.

### scRNA-seq

To profile NRF1-driven transcriptomic changes, NRF1 mRNA-transfected MSCs, SCR mRNA-transfected MSCs, and NT MSCs were processed for scRNA-seq by EMPIRI Inc. (Houston, TX). High-viability cell suspensions (>95%) were loaded onto the 10x Chromium Controller using ChipK, aiming for a capture rate of 5,000 cells per condition. The resulting single-cell cDNA libraries were generated using the Chromium Next GEM Single Cell 5’ Reagent Kits v2 (10x Genomics) according to standard protocols.

### scRNA-seq analysis

Raw Illumina sequencing reads were aligned to the reference genome GRCh38-2024-A for Homo Sapiens using cellranger-9.1.0 in 10x Genomics Cloud Analysis. This study includes three Fastq datasets: one for NRF1 overexpression, and the others for NT and SCR mRNA transfected MCS respectively. Subsequent analysis of these datasets utilized Seurat v.5.3.0. We excluded low-quality and doublet cells by keeping cells with nFeatures between 200 and 9,500, and ratio of mitochondrial genes smaller than 5%. Each dataset was independently normalized relative to library size, log_2_-transformed, scaled and carried out principal component analysis. The three datasets were then integrated using reciprocal PCA (RPCA) following the standard protocol in Seurat [37, 38], producing an integrated single-cell dataset with gene expression counts across 34,371 unique features for 13,628 cells.

Clustering used a Louvain graph-based approach with resolution 0.5, identifying 14 clusters. Cluster specific marker genes were identified with Wilcoxon rank-sum tests using wilcoxauc function in R package presto, applying thresholds logFC >0.25, min.pct >0.1, and adjusted p <0.1. Differential expression between NRF1 and CTR cells within specified clusters was also computed with wilcoxauc, and recorded expression profile for all the genes expressed in more than 10% of NRF1 cells. GSEA was performed using the pseudo bulk DEG (differentially expressed genes) profile described above. For clusters where more than 90% (C4, 5 and 11) or less than 10% (C0, 3, 8, 10, 12 and 13) of cells were from CTR, DEGs comparing this cluster vs. all others were taken, while for others (C1, 2, 6, 7 and 9), DEGs from NRF1 vs CTR were taken, while also included this comparison within C0. A -log10(p-value)*sign(logFC) was used as the score for each gene in the comparison, and applied preranked GSEA using GSEA v4.4.0 to test 12 GO terms and 1 KEGG pathways. The list of member genes was obtained from Human MSigDB v2026.1.HS.

### Oxidative stress induction in ECs

EC oxidative injury was induced by treatment with 250 µM H_2_O_2_ for 1 h. Cells were then washed twice with PBS to remove residual oxidant prior to co-culture.

### Co-culture conditions

Immediately after EC washing and 6 h post-transfection of MSCs, co-culture experiments were initiated on 6-well plates. For direct co-culture assays, MSCs and ECs were mixed at a 1:3 ratio. Considering the larger somatic size of MSCs compared to ECs, a total of 1.2 × 10^5^ cells were seeded per well. The combined cell suspension was carefully plated to ensure uniform distribution and prevent cell aggregation. For indirect co-culture assays to evaluate endothelial function, a Boyden chamber system (6-well Transwell ThinCert™ inserts, 8.0 µm pore size) was employed. ECs were seeded in the lower chamber at a density of 1.5 × 10^5^ cells/well. Transfected MSCs were seeded in the upper chamber at a density of 0.5 × 10^5^ cells/well, maintaining a 1:3 ratio of MSCs to ECs. The cells were co-cultured for 18 h. Upon completion of the incubation period, the experiment was terminated, and cells were processed for downstream analyses.

### Assessment of mitochondrial transfer of mitochondria from MSCs to ECs

MSCs with AcGFP1-labeled mitochondria (donors) transfected with SCR or NRF1 mRNA were co-cultured with ECs with mito-mCherry-labeled mitochondria (recipients) under direct and transwell conditions. For imaging, co-cultures were washed with PBS and fixed (4% PFA). To visualize the cytoskeletal and TNT architecture, cells were permeabilized with 0.1% Triton X-100 in PBS for 5 min. Subsequently, cells were stained with deep red-conjugated phalloidin for 30 min to label F-actin, followed by nuclear counterstaining with DAPI for 15 min at room temperature at dark. The slides were then mounted for imaging. Mitochondrial transfer was visualized by identifying MSC-derived AcGFP1 signals colocalized within mCherry-positive ECs.

For quantitative analysis via flow cytometry, cells were trypsinized after co-culture, washed, and resuspended in FACS buffer. ECs were identified by gating on the mito-mCherry positive population. Subsequently, the mean fluorescence intensity (MFI) of the AcGFP1 signal within this gated population was analyzed to quantify mitochondrial transfer efficiency.

To investigate the intercellular transfer of mitochondria and their subsequent colocalization with the host mitochondrial network, imaging was performed for indirect co-culture conditions. The degree of colocalization between the MSC-derived mitochondria and EC mitochondria was quantified using Mander’s Coefficiency test via the Coloc2 plugin in Fiji software.

### Quantification of heterotypic intercellular connections between MSCs and ECs

To quantify TNT-mediated donor-recipient connectivity, direct co-cultures of mito-AcGFP1 MSCs and mito-mCherry ECs were imaged by confocal microscopy. Heterotypic TNTs connecting an AcGFP1⁺ donor MSC to an mCherry⁺ recipient EC were manually scored using the ICY bioimage analysis software across ten random fields per well. Homotypic connections (same fluorophore) and non-bridging protrusions were excluded. Heterotypic TNT abundance was quantified by dividing the number of heterotypic TNTs by the total number of interconnected MSCs and ECs in the field.

### MitoSOX assay

Mitochondrial superoxide production was assessed. Following mRNA transfection and indirect co-culture, ECs cultured on collagen-coated coverslips were incubated with MitoSOX™ Red for 15 min, fixed with 4% PFA, and counterstained with DAPI. Fluorescence images were acquired using a Nikon A1 confocal microscope. Quantitative analysis was performed using an LSRII flow cytometer by measuring mean fluorescence intensity.

### JC-1 assay

Mitochondrial membrane potential was assessed using JC-1 assay. Following mRNA transfection and indirect co-culture, ECs cultured on collagen-coated coverslips were incubated with JC-1 dyes for 15 min, fixed with 4% PFA, and counterstained with DAPI. Fluorescence images were acquired using a Nikon A1 confocal microscope. Quantitative analysis was performed using an LSRII flow cytometer by measuring mean or median fluorescence intensity. The ratio of red (J-aggregate) to green (monomer) fluorescence was calculated as an indicator of mitochondrial membrane polarization.

### Seahorse bioenergetics and ATP quantification

For measurement of cell respiration, harvested ECs were seeded into XF96 cell culture microplates (Agilent Technologies, Santa Clara, CA, USA) at a density of 2 × 10^4^ cells/well and allowed to attach overnight. The oxygen consumption rate (OCR) and extracellular acidification rate (ECAR) were measured using an Agilent Seahorse XFe96 Extracellular Flux Analyzer. On the day of the assay, the culture medium was replaced with XF Base Medium (pH 7.4) supplemented with 10 mM glucose, 1 mM sodium pyruvate, and 2 mM L-glutamine. The cells were incubated in a non-CO_2_ incubator at 37°C for 1 h to equilibrate the medium. During the assay, mitochondrial modulators were sequentially injected at the following final concentrations: 1.5 µM oligomycin (ATP synthase inhibitor), 1.0 µM FCCP (mitochondrial uncoupler), and a mixture of 0.5 µM rotenone and antimycin A (Complex I and III inhibitors). OCR and ECAR values were normalized to the total cell number (per 1,000 cells), determined by nuclei counting following the assay. Data analysis was performed using Wave Controller Software (Agilent Technologies).

Intracellular ATP levels were quantified using the ATPlite Luminescence Assay System (Revvity, Waltham, MA, USA). Harvested ECs were seeded into white opaque 96-well plates at a density of 1 × 10^4^ cells/well and cultured overnight. To induce cell lysis, 50 µl of mammalian cell lysis solution was added to each well, and the plates were agitated on a shaker for 5 min at 700 rpm. Subsequently, 50 µl of substrate solution was added, followed by an additional 5 min of shaking at 700 rpm. Plates were then incubated for 10 min at room temperature in the dark to stabilize the luminescent signal. ATP-dependent luminescence was measured using a microplate reader, and the ATP levels were reported as counts per second (CPS).

### Senescence-associated β galactosidase (SA-β-gal) assay

To assess SA-β-gal activity, ECs were washed with PBS and fixed with a fixation solution composed of 2% formaldehyde (vol/vol) and 0.2% glutaraldehyde (vol/vol) in PBS for 5 min. Following fixation, cells were washed and incubated for 8 h at 37°C in a CO_2_ free incubator while immersed in a prepared X-gal staining solution. The staining solution consisted of X-gal (Invitrogen, Waltham, MA), 40 mM citric acid/sodium phosphate buffer (pH 6.0), 5 mM potassium hexacyanoferrate (II) trihydrate, 5 mM potassium hexacyanoferrate (III), 150 mM NaCl, and 2 mM. After staining, ECs were washed with PBS, and images were acquired using the EVOS cell imaging system. SA-β-gal-positive cells were manually counted and expressed as a percentage of the total cell number.

### Apoptosis by Annexin V/PI

To evaluate the protective effects of NRF1 primed MSCs against oxidative stress-induced apoptosis, ECs were exposed to 250 µM H_2_O_2_ for 1 h and subsequently subjected to indirect co-culture with SCR- or NRF1-transfected MSCs. After the co-culture, ECs were harvested and washed twice with cold PBS. The cells were then resuspended in binding buffer and stained with Annexin V-FITC and Propidium Iodide (PI) for 15 min at room temperature in the dark. Apoptotic cells were analyzed using flow cytometry. The total percentage of apoptotic cells was calculated by the populations of early apoptotic (Annexin V^+^/PI^-^) and late apoptotic (Annexin V^+^/PI^+^) cells. The quantitative data were presented as bar graphs.

### Tube formation assay

To determine the angiogenic potential of endothelial cells, a tube formation assay was performed. Matrigel (Corning, Glendale, AZ) was thawed on ice to prevent premature polymerization. The cold Matrigel was carefully dispensed into a 96 well plate (50 µl/well) on ice, and the plate was incubated at 37°C for 30 min to allow the matrix to solidify. Subsequently, ECs were seeded at a density of 1 × 10^4^ cells/well onto the polymerized Matrigel and cultured in MSC culture medium. After 24 h of incubation, capillary-like tube structures were visualized using the EVOS imaging system. The degree of angiogenesis was quantified by measuring the total number of branching points using Fiji software as published previously [39].

### eNOS analysis

ECs were exposed to 250 µM H_2_O_2_ for 1 h and subsequently co-cultured with SCR- or NRF1-transfected MSCs. Following co-culture, ECs were washed with PBS and incubated in serum-free EC culture medium (0% FBS) for 6 h to induce starvation. To stimulate eNOS activity, the cells were treated with 100 nM Bradykinin for 30 min. Total protein was extracted using a RIPA buffer (GenDEPOT, Houston, TX) supplemented with a phosphatase and protease inhibitor cocktail (GenDEPOT). The levels of phosphorylated eNOS (Ser1177) and total eNOS were assessed by Western blot analysis.

### Western blot

Protein expression was analyzed by Western blot. Prior to cell lysis, cells were washed with cold PBS and supplemented with a phosphatase and protease inhibitor cocktail. Cells were lysed using RIPA buffer on ice, and lysates were cleared by centrifugation. Protein concentrations were determined using a DC Protein Assay kit (Bio-Rad, Hercules, CA). To extract total protein from EVs, pellets derived from MSC supernatant were lysed in 30 µL of RIPA buffer supplemented with protease and phosphatase inhibitor cocktails. To ensure complete lysis and release of intra-vesicular proteins, the samples were vortexed three times for 25 s each. Subsequently, the lysates were centrifuged at 24,000 × g for 30 min at 4°C to remove insoluble debris, and the supernatants were collected for Western blot analysis. Equal amounts of protein were loaded onto precast SDS–PAGE gels (Bio-Rad) and separated under denaturing conditions, followed by transfer onto nitrocellulose membranes (Bio-Rad). Membranes were blocked in 5% BSA prepared in Tris-buffered saline containing 0.1% Tween-20 (TBST) and incubated overnight at 4 °C with the appropriate primary antibodies. Primary antibodies included: NRF1 (Abcam, 1:1000), TFAM (Cell Signaling Technology, 1:1000), COXIV (Cell Signaling Technology, 1:5000), Talin2 (abcam, 1:1000), FAK (Cell Signaling Technology, 1:1000), Cdc42 (Cell Signaling Technology, 1:1000), Miro1 (Abcam, 1:1000), EPS8 (Cell Signaling Technology, 1:1000), CD63 (abcam, 1:1000), HIF-1α (Cell Signaling Technology, 1:1000), HKII (Cell Signaling Technology, 1:1000), PFKFB3 (Cell Signaling Technology, 1:1000), LDH-B (Abcam, 1:1000), DRP1 (Cell Signaling Technology, 1:5000), MFN2 (Cell Signaling Technology, 1:5000), OPA1 (Abcam, 1:5000), FIS1 (Santa Cruz Biotechnology, 1:2000), PINK1 (Cell Signaling Technology, 1:1000), Parkin (Cell Signaling Technology, 1:1000), LC3B (Cell Signaling Technology, 1:1000), p53 (Santa Cruz Biotechnology, 1:1000), p21 (Abcam, 1:500), p16 (Abcam, 1:500), Bcl-2 (Cell Signaling Technology, 1:1000), BAX (Cell Signaling Technology, 1:1000), ICAM-1 (Cell Signaling, 1:1000), VCAM-1 (Cell Signaling, 1:100), COX2 (SantaCruz, 1:500), VEGF-A (Cell Signaling, 1:1000), peNOS (Cell signaling Technology, 1:1000), eNOS (Cell Signaling Technology, 1:1000) and β-actin (Cell Signaling Technology, 1:10,000) as a loading control.

After primary antibody incubation, membranes were washed three times with TBST and incubated with HRP-conjugated secondary antibodies (anti-rabbit or anti-mouse IgG; Cell Signaling Technology, 1:10,000) for 2 h at room temperature. Membranes were washed again with TBST and protein bands were visualized using enhanced chemiluminescence substrate (Immobilon HRP, Millipore Sigma). Signals were detected using a ChemiDoc imaging system (Bio-Rad), and band intensities were quantified by densitometric analysis.

### Statistical Analysis

All data are presented as mean ± standard error of the mean (SEM). Statistical analyses were performed using GraphPad Prism software. Unless otherwise stated, all experiments were performed with at least three biological replicates (n ≥ 3). For imaging-based quantifications, ten random fields per well were analyzed. Unless otherwise indicated, comparisons between groups were conducted using one-way ANOVA followed by Dunnett’s multiple comparison test. Statistical significance was set up at p < 0.05.

## Results

### NRF1-driven mitochondrial biogenesis couples to cytoskeletal remodeling and increased TNT formation

NRF1 induction increased mitochondrial mass in MSCs, reflected by greater mitochondrial-associated AcGFP1 fluorescence (Supplementary Fig. S1a, b). NRF1, TFAM, and COXIV, the latter two canonical biogenesis readouts, with TFAM supporting mtDNA transcription/replication, and COXIV widely used as a mitochondrial marker [40, 41], were elevated in NRF1-overexpressing MSCs compared with CTR and SCR controls (Supplementary Fig. S1c), consistent with our previous findings [32].

To determine whether NRF1 also programs a pro-transfer cytoskeletal state, we profiled regulators of actin remodeling and focal-adhesion assembly. Talin2, FAK, and Cdc42 were profiled because Rho GTPase-driven actin remodeling [42] and focal-adhesion assembly are key prerequisites for membrane protrusions/TNT-like structures [43, 44]. Immunoblotting revealed significant upregulation of Talin2 (∼3-fold change), FAK (∼2-fold change), and Cdc42 (∼3-fold change) in NRF1-overexpressing MSCs versus SCR controls (Fig. 1a). NRF1 also increased Miro1 (∼2-fold change) and EPS8 (∼1.4-fold change) compared to SCR controls (Fig. 1b), linking actin dynamics to mitochondrial trafficking and TNT formation [45–47]. Confocal imaging showed an increase in TNT-like structures in NRF1 MSCs relative to SCR controls (Fig. 1c; Supplementary Fig. S2).

**Figure 1.**
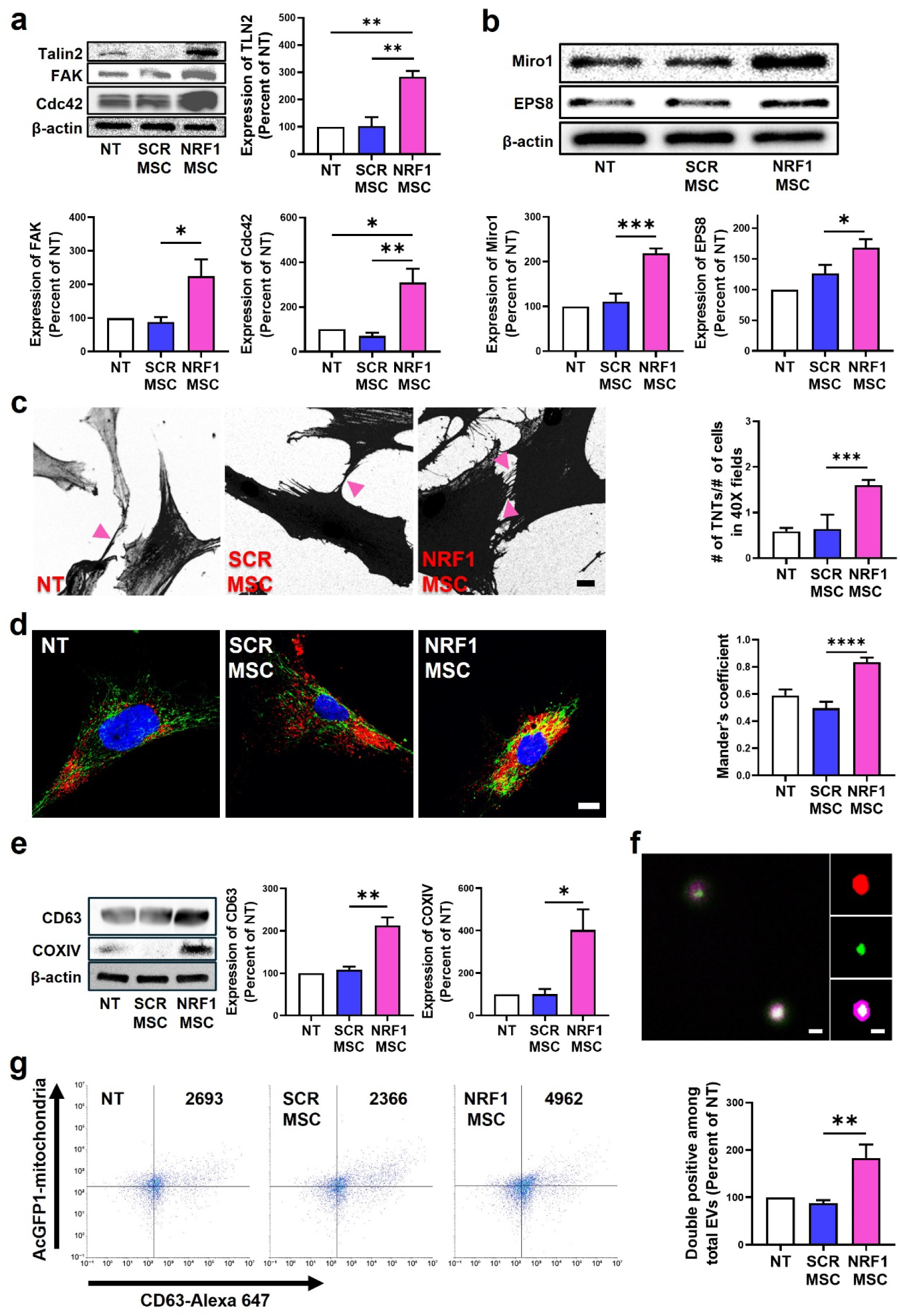
NRF1 priming engages mitochondrial transfer machinery, resulting in cytoskeletal reorganization, TNT formation, and increased release of mitochondria-containing EVs. MSCs were transfected with either scrambled (SCR) or NRF1 mRNA. Controls consisted of non-transfected MSCs (NT). a) Representative western blot and densitometric analysis of expression of the focal adhesion and cytoskeletal proteins Talin2, FAK, and Cdc42. b) Representative western blot and densitometric analysis of Miro1 and EPS8 protein expression. c) Representative fluorescence images showing MSC TNT formation (indicated by pink arrowheads). Images are represented monochromatic (black and white) for clarity of presentation. Original fluorescence images available in Supplementary Materials (Supplementary Materials Fig. S2). Scale bar = 10 µm. The bar graph represents the number of TNTs per the number of cells per 40x field. d) Representative confocal microscopy images showing subcellular distribution and colocalization of mitochondria (green) and CD63 (red). Nuclei were stained with DAPI (blue). Scale bar = 10 µm. Quantitative analysis of the colocalization between mitochondria-and CD63-associated fluorescence using Mander’s overlap coefficient. e) Representative western blot and densitometric analysis of CD63 and COXIV in MSC-derived EVs. f) Confocal microscopy images of EVs labeled for mitochondrial content (AcGFP1) and the EV surface marker CD63 (red), with images to the right highlighting individual channels. Scale bar = 1 µm. g) Representative scatter plots of populations of MSC-derived EVs analyzed for MSC mitochondria-associated AcGFP1 and the EV marker CD63 (Alexa Fluor 647). Values in the upper right quadrant indicate absolute count of double-positive EVs (AcGFP1^+^/CD63^+^), with the number of mitochondria-containing EVs among total EVs quantified in the accompanying graph. For all Western blots, β-actin was used as a loading control. For densitometric analysis, protein markers were normalized to β-actin expression levels relative to NT. Data are presented as mean ± SEM. Statistical significance is indicated as \**p* < 0.05, \*\**p* < 0.01, \*\*\**p* < 0.001, and \*\*\*\**p* < 0.0001 vs CTR or SCR.

### NRF1 increases mitochondrial association with CD63^+^ compartments and enhances release of mitochondria-loaded EVs

To evaluate whether NRF1 induction promotes EV-mediated mitochondrial export, we quantified co-localization between AcGFP1-labeled mitochondria and CD63, a marker enriched in endosomal/exosomal membranes that reports engagement of MVB/EV trafficking routes [48]. NRF1 MSCs showed increased co-localization between AcGFP1-labeled mitochondria and CD63-positive compartments in NRF1-primed MSCs (Fig. 1d), confirmed by higher Mander’s overlap coefficients.

Nanoparticle tracking analysis of MSC supernatants showed NRF1 MSCs released more EVs than controls (Supplementary Fig. S3), without major shifts in EV size distributions (Supplementary Fig. S4). EV immunoblotting demonstrated increased COXIV alongside CD63 in NRF1 MSC isolates (Fig. 1e). Super-resolution confocal microscopy imaging identified AcGFP1^+^ mitochondrial signal associated with CD63^+^ EV compartments (Fig. 1f). NanoFlow cytometry further showed a nearly 1-fold increase in AcGFP1^+^/CD63^+^ particles in the NRF1-primed MSC group compared to both NT and SCR controls (Fig. 1g). Together, these data show that NRF1 overexpression enhances both TNT formation and secretion of mitochondria-loaded EVs.

### scRNA-seq profiling supports NRF1-associated activation of TNT and vesicle trafficking programs

scRNA-seq analysis was used to elucidate the impact of NRF1 overexpression on coordinated, pathway-level transcriptional programs involving cytoskeletal/adhesion and vesicle-trafficking modules. Transcriptional profiles of NRF1 mRNA overexpressed cells were compared against a control group (CTR) that combined NT and SCR mRNA transfected MSCs due to their high similarity in gene expression patterns. Uniform Manifold Approximation and Projection (UMAP) visualization showed a large shift in population distribution between the two groups (Fig. 2a). While CTR cells clustered densely in specific regions (predominantly Clusters 0, 1, 2, and 3), NRF1 overexpressing cells showed distinct and separated clusters. Quantitative analysis of cell origin supported this observation. Notably, Cluster 4 (C4) and Cluster 5 (C5) were almost exclusively composed of NRF1-overexpressed cells (1,189 and 1,135 cells in NRF1 vs. 34 and 28 in CTR, respectively), whereas Cluster 0 was predominantly occupied by control cells (Fig. 2a). This suggests that NRF1 induction drives MSCs into specific, novel transcriptional states.

**Figure 2.**
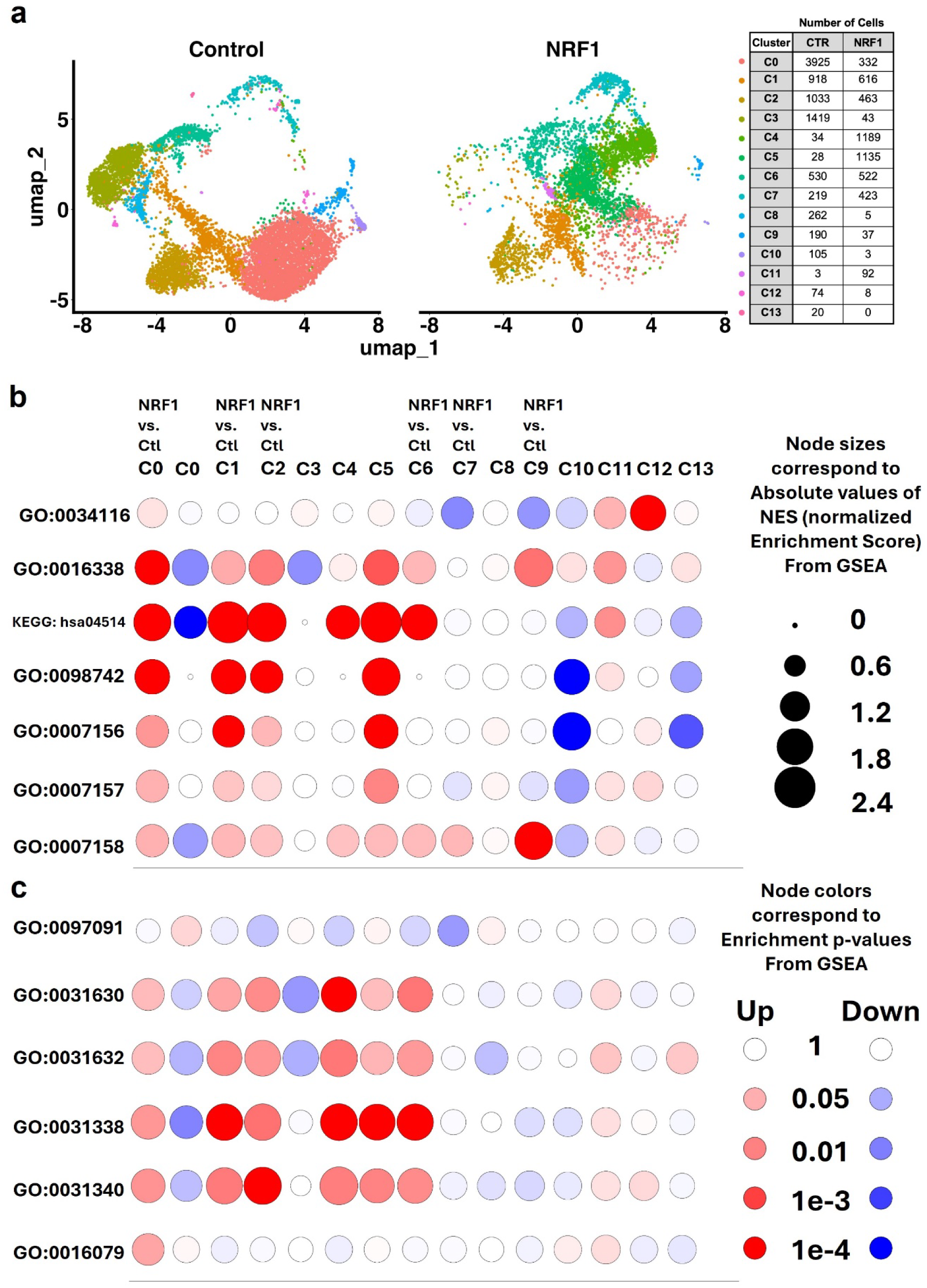
scRNA-seq profiling reveals NRF1-driven reprogramming toward adhesive and secretory states. MSCs were transfected with either scrambled (SCR) or NRF1 mRNA. Controls consisted of non-transfected MSCs (NT). a) UMAP visualization comparing NRF1-primed MSCs against the control (CTR) group that combines NT and SCR MSCs. The inset table quantifies the population shift (number of MSCs in each cluster). For bubble charts in b and c, GSEA Gene set IDs can be found in Table 1. b) Bubble chart demonstrating GSEA in different clusters (C0-C13) illustrating activation of specific pathways specific to adhesion molecule pathways. c) Bubble chart demonstrating GSEA in different clusters (C0-C13) illustrating activation of pathways specific to intracellular vesicle dynamics such as formation, fusion, and trafficking. For specific clusters where NRF1 cell numbers were insufficient, NRF1-derived cells within the cluster were compared against the total CTR population (displays as Cn vs CTR). Node size represents the absolute Normalized Enrichment Score (NES), while color indicates statistical significance and regulation direction (Red: Upregulated; Blue: Downregulated).

**Table 1.**
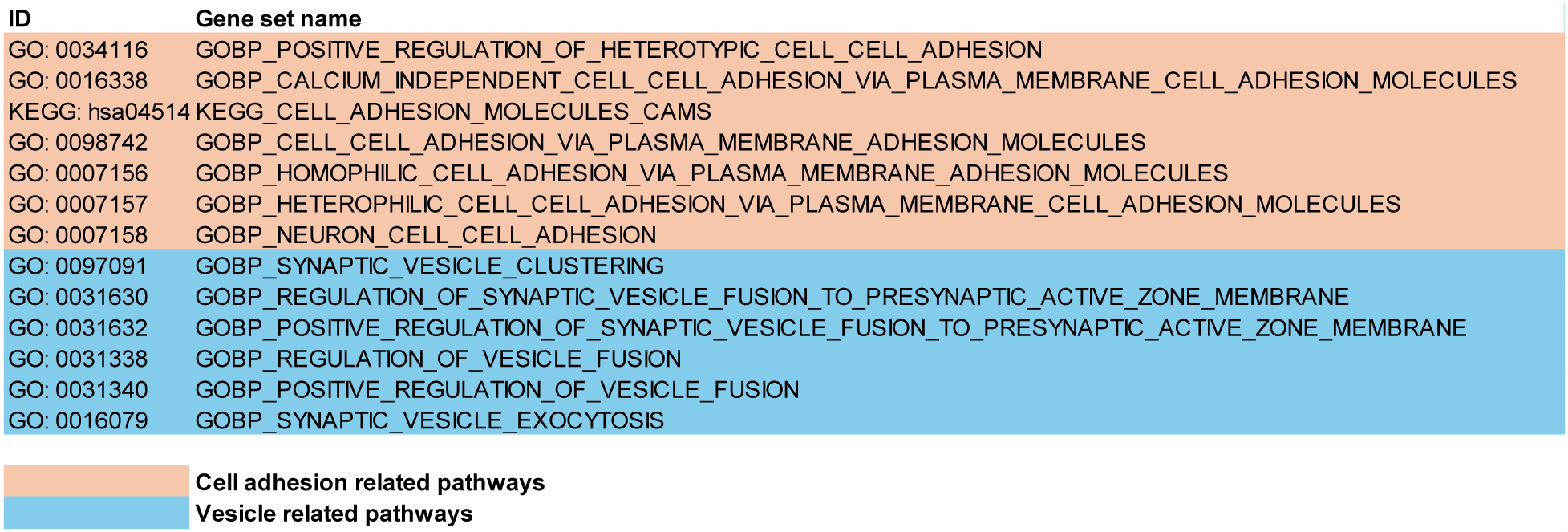
GSEA Gene set IDs.

Gene Set Enrichment Analysis (GSEA) was then performed across all clusters to examine cell-cell interaction capabilities and secretory activity (Table 1, Fig. 2b). Bubble chart analysis revealed a striking upregulation of adhesion-related pathways in NRF1-enriched clusters (C4 and C5). Specifically, gene sets for “Cell adhesion molecules (CAMs)” and “Cell-cell adhesion via plasma membrane adhesion molecules” were significantly enriched in C4 and C5. Conversely these were non-significant or downregulated in CTR dominant clusters (C1 and C10). Representative GSEA enrichment plots (Supplementary Materials Fig. S5) validated these findings, showing strong positive enrichment scores for KEGG Cell Adhesion Molecules pathways in both C4 and C5. In addition, FN1 and SSPN were identified as the top upregulated cell adhesion markers in C4 and C5 (Supplementary Materials Fig. S6). Furthermore, a broader GSEA analysis across different MSC subpopulations revealed a diverse landscape of enriched cell adhesion pathways (Supplementary Materials Fig. S7). Notably, the KEGG Cell Adhesion Molecules pathways was consistently enriched across multiple clusters (C1, C2, C3, and C6), which is notable given that, despite not having a particularly pronounced presence of NRF1-derived cells, still exhibited a balanced proportion or substantial presence of NRF1-derived cells. Other distinct adhesion pathways, such as calcium-independent or neuron cell-cell adhesion, were identified in the remaining subpopulations. Concurrently, the analysis for secretory activity highlighted a robust upregulation of vesicle dynamics in the NRF1-dominant populations (Fig. 2c). Both C4 and C5 displayed significant positive enrichment for Regulation of vesicle fusion, Positive regulation of vesicle fusion, and Synaptic vesicle fusion to presynaptic active zone membrane. This functional signature was largely absent or suppressed in CTR clusters (C0, C1, C2). GSEA enrichment plots (Supplementary Materials Fig. S8) further corroborated these results, demonstrating a clear positive correlation between NRF1 overexpression and the regulation of vesicle fusion in both C4 and C5. CD63 and CLU were shown as the most significant markers related to vesicle dynamics in the C4 and C5 (Supplementary Materials Fig. S9). Furthermore, GSEA analysis confirmed that pathways regulating vesicle fusion were consistently enriched across multiple MSC subpopulations (C1, C2, C4, C5, and C6), suggesting a broad impact of NRF1 on vesicle dynamics. Specifically, in clusters C1, C2, and C6, where NRF1-derived cells do not appear as the major population, using a direct comparison between NRF1 and whole CTR groups revealed a robust enrichment of these vesicle fusion pathways (Supplementary Materials Fig. S10). Taken together, these data demonstrate that NRF1 overexpression fundamentally reprograms MSCs, driving them into distinct cell states (C4, C5) characterized by enhanced structural connectivity (adhesion) and active vesicle fusion machinery.

### NRF1 enhances mitochondrial transfer to oxidatively stressed ECs via contact-dependent and contact-independent routes

To determine whether NRF1 priming increases mitochondrial donation, ECs exposed to oxidative stress were co-cultured with MSCs either directly or using transwell setups to separate contact-dependent from soluble/EV-mediated transfer. In direct co-culture, confocal microscopy revealed TNT-like F-actin bridges between NRF1 MSCs and stressed ECs (Fig. 3a; Supplementary Fig. S11), with more prominent formation of intercellular connections in the NRF1 MSC group compared to SCR controls (Supplementary Fig. S12). Notably, AcGFP1-labeled mitochondria were visible within these bridges and within recipient ECs (Fig. 3a; Supplementary Fig. S12). Flow cytometry showed increased AcGFP1 MFI in recipient ECs in the NRF1 condition (Fig. 3a). In transwell assays, AcGFP1 signal was detected in stressed ECs and overlapped with the endogenous mitochondrial network (Fig. 3b; Supplementary Fig. S13). Recipient AcGFP1 MFI also increased in the NRF1 condition versus SCR (Fig. 3b). Taken together, NRF1 induction increases mitochondrial transfer through both contact-dependent and contact-independent mechanisms.

**Figure 3.**
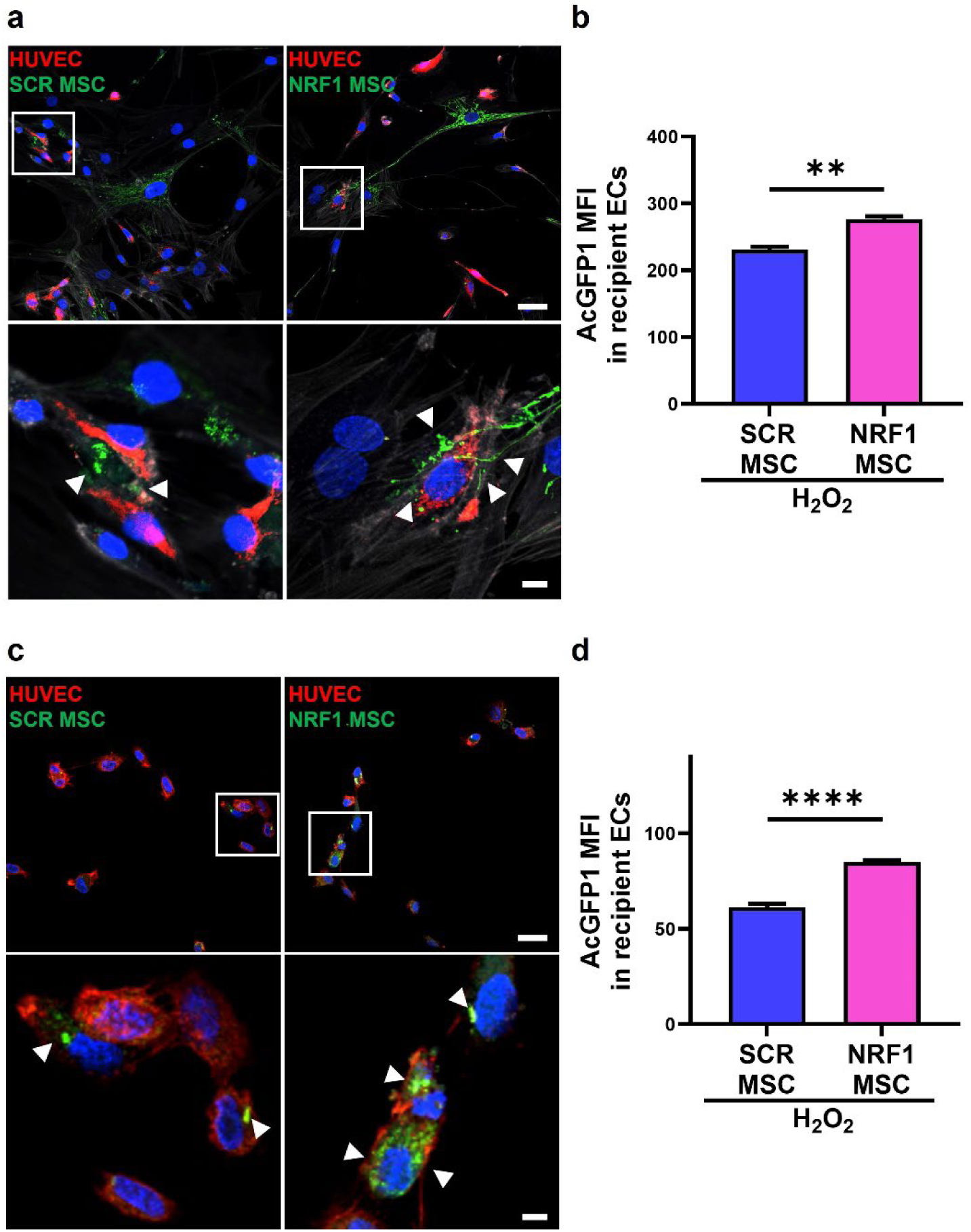
NRF1 priming of MSCs enhances mitochondrial delivery to ECs undergoing oxidative stress. H_2_O_2_-exposed (250 µM, 1 h) ECs were co-cultured with MSCs either directly or indirectly using transwell setups to separate contact-dependent from soluble/EV-mediated transfer. MSCs were transfected with either scrambled (SCR) or NRF1 mRNA. Lentiviral transduction was performed to obtain MSCs with mitochondria expressing AcGFP1 (green) and ECs with mitochondria expressing mCherry (red). In all images, cell cytoskeleton was stained with F-actin (gray) and nuclei with DAPI (blue). a) Representative confocal microscopy images of ECs co-cultured with SCR- or NRF1-transfected MSCs in a direct contact co-culture system. White arrowheads indicate AcGFP1-labeled mitochondria transferred to ECs. Scale bar = 25 µm (upper image) and 5 µm (lower image). b) Mean fluorescence intensity (MFI) of AcGFP1 in recipient ECs in the direct co-culture system, as determined by flow cytometry. c) Representative confocal microscopy images of ECs co-cultured with SCR- or NRF1-transfected MSCs in an indirect transwell co-culture system. Scale bar = 25 µm (upper image) or 5 µm (lower image). d) MFI of AcGFP1 in recipient ECs in the indirect co-culture system, as determined by flow cytometry. Data are presented as mean ± SEM. *\*\*p*<0.01, *\*\*\*\*p*<0.0001 vs SCR.

### NRF1-primed MSCs reduce mtROS and preserve mitochondrial membrane potential in stressed ECs

Oxidative injury disrupts respiratory chain function and depolarizes mitochondrial membrane potential (ΔΨm) [44]. Herein, H_2_O_2_ increased mitochondrial superoxide (MitoSOX) (Fig. 4a, b), while NRF1 MSC co-culture reduced MitoSOX MFI by ∼25% versus SCR (Fig. 4a, b). In JC-1 assays, H_2_O_2_ decreased the J-aggregate signal and increased monomer signal, consistent with ΔΨm depolarization (Fig. 4c, d). NRF1 MSC co-culture increased the J-aggregate/J-monomer ratio by ∼2-fold change versus SCR (Fig. 4c, d), indicating ΔΨm preservation.

**Figure 4.**
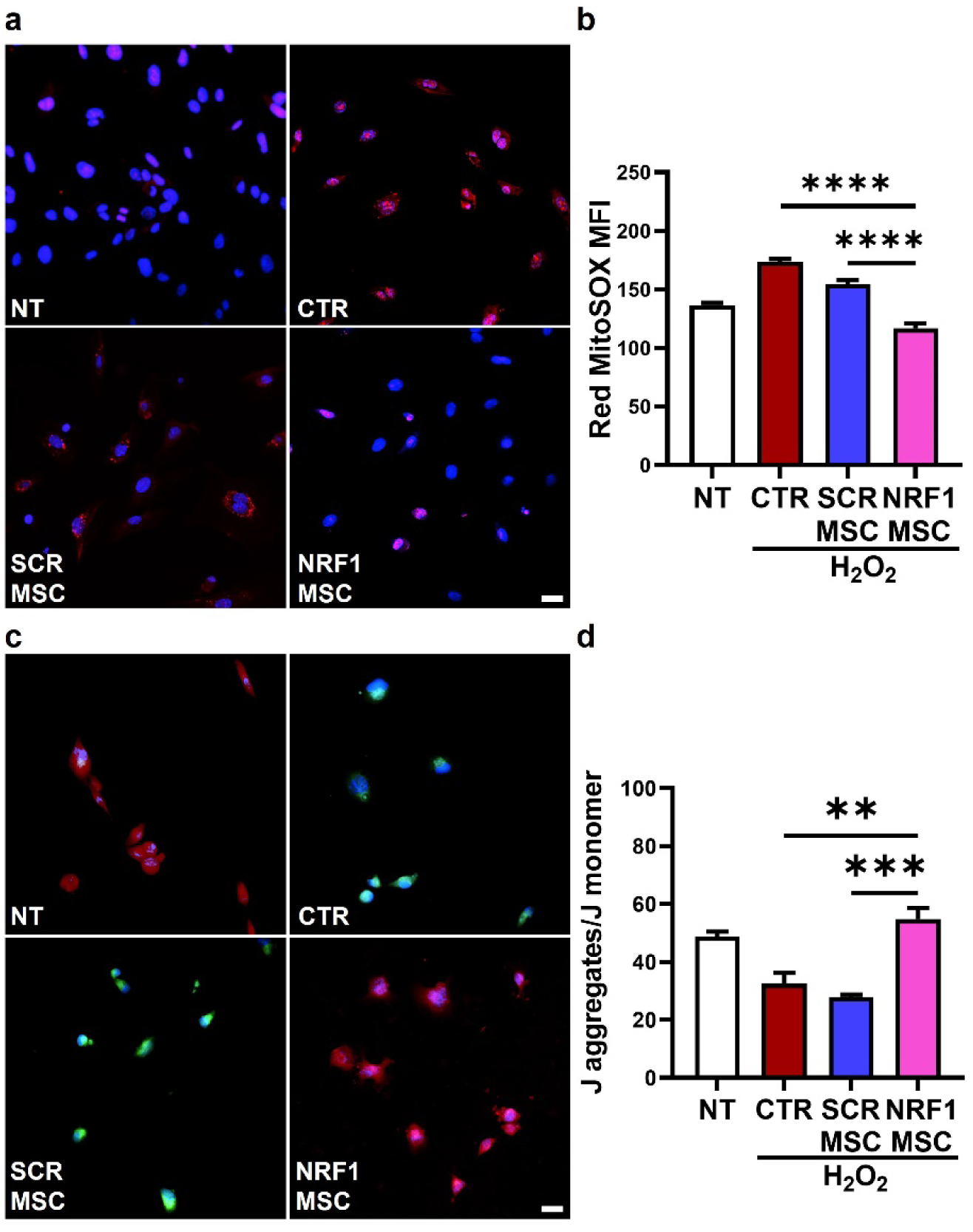
NRF1-primed MSCs alleviate mitochondrial oxidative stress and preserve mitochondrial membrane potential in ECs. H_2_O_2_-exposed (250 µM, 1 h) ECs were indirectly co-cultured with MSCs transfected with either scrambled (SCR) or NRF1 mRNA. Controls consisted of healthy, non-treated ECs (NT) and H2O_2_-exposed ECs indirectly co-cultured with non-transfected MSCs (CTR). a) Representative confocal microscopy images depicting MitoSOX stained MSCs. Oxidized MitoSOX reagent is represented in red and DAPI-stained nuclei appear in blue. Scale bar = 25 µm. b) Mean fluorescence intensity (MFI) of oxidized MitoSOX reagent quantified by flow cytometry. c) Representative confocal microscopy images of MSCs undergoing JC-1 assay to assess mitochondrial membrane potential. JC-1 monomers are represented in green, J aggregates in red, and DAPI-stained nuclei in blue. Scale bar = 25 µm. d) The ratio of the median fluorescence intensity (MedFI) of J aggregates to JC-1 monomers quantified by flow cytometry. Data are presented as mean ± SEM. *\*\*p*<0.01, *\*\*\*p*<0.001, *\*\*\*\*p*<0.0001 vs H_2_O_2_ or SCR.

### NRF1-primed MSCs improve bioenergetics and suppress stress-associated glycolytic reprogramming

Extracellular flux analysis showed that H_2_O_2_ suppressed OCR and increased ECAR, lowering the OCR/ECAR ratio to ∼0.4 versus ∼0.6 in NT controls (Fig. 5a). NRF1 MSC co-culture restored the OCR/ECAR ratio toward NT (Fig. 5a). ATP decreased to ∼50% of NT after H_2_O_2_. Consistent with improved respiration, ATP increased by ∼40% with respect to SCR following NRF1 MSC co-culture (Fig. 5b). Given the link between oxidative stress and glycolytic reprogramming [49], we assessed HIF-1α and glycolytic enzymes (HKII, PFKFB3, LDH-B). H_2_O_2_ increased HIF-1α (∼2-fold change vs NT) and induced HKII, PFKFB3, and LDH-B, whereas NRF1-primed MSC co-culture yielded a ∼2-fold decrease in HIF-1α compared to CTR and suppressed glycolytic enzyme induction (Fig. 5c).

**Figure 5.**
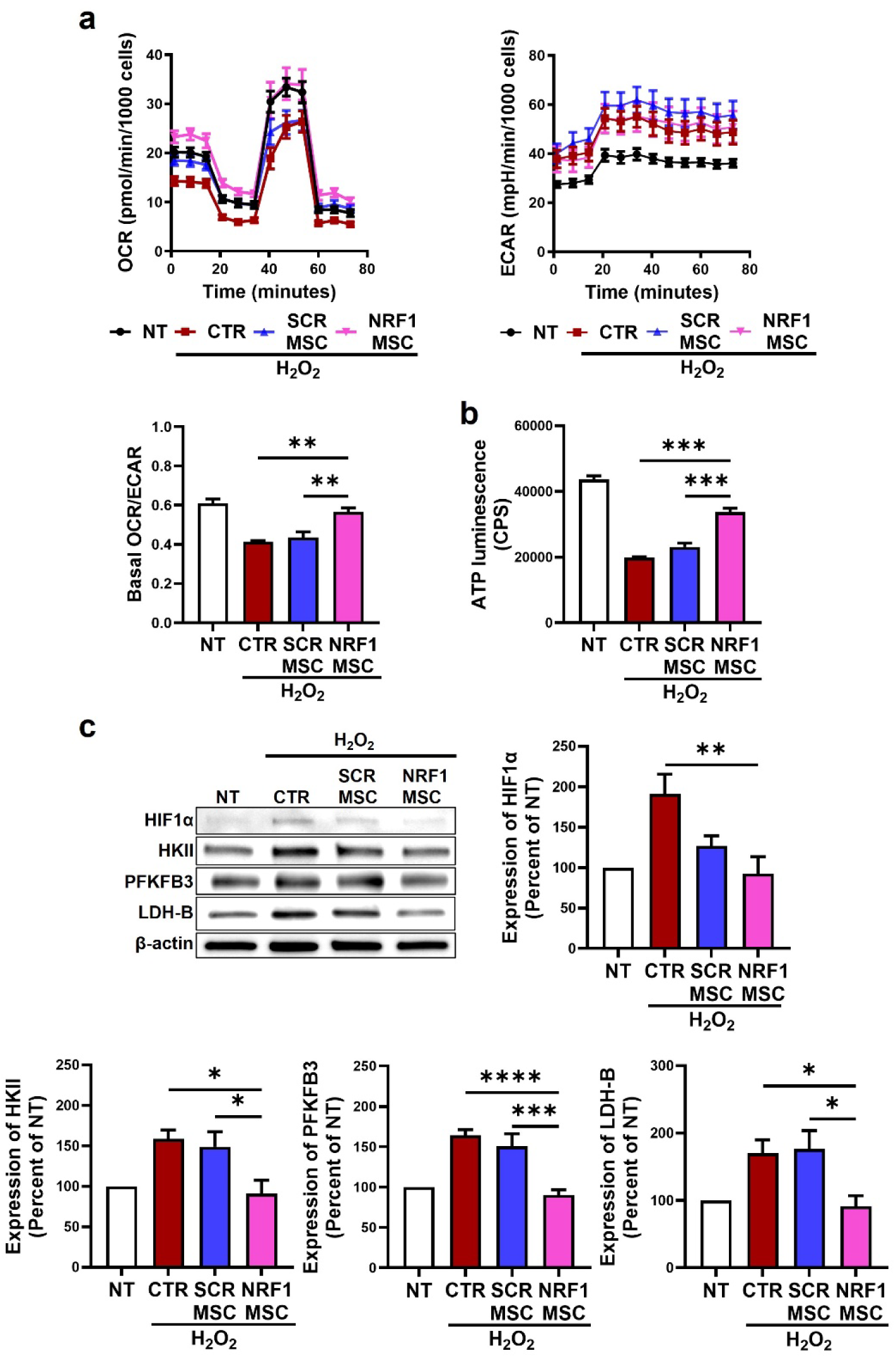
NRF1-primed MSCs improve mitochondrial bioenergetics in ECs following oxidative stress. H_2_O_2_-exposed (250 µM, 1 h) ECs were indirectly co-cultured with MSCs transfected with either scrambled (SCR) or NRF1 mRNA. Controls consisted of healthy, non-treated ECs (NT) and H_2_O_2_-exposed ECs indirectly co-cultured with non-transfected MSCs (CTR). a) Bioenergetic analysis of ECs via examination of the oxygen consumption rate (OCR) and extracellular acidification rate (ECAR). O: oligomycin; F: FCCP; R + A: rotenone + antimycin A. The bar graph represents the ratio of basal OCR/ECAR of ECs. b) Relative intracellular ATP of ECs. c) Representative western blot of glycolytic regulator enzymes (HIF1ɑ, HKII, PFKFB3, LDH-B) in ECs. β-actin was used as a loading control. For densitometric analysis, protein markers were normalized to β-actin expression levels relative to NT. Data are presented as mean ± SEM. *\*p*<0.05, *\*\*p*<0.01, *\*\*\*p*<0.001, *\*\*\*\*p*<0.0001 vs H_2_O_2_ or SCR.

### NRF1-primed MSCs rebalance mitochondrial dynamics and restore mitophagy-associated signaling in stressed ECs

We then examined the effect of NRF1-primed MSC mitochondrial transfer on mitochondrial dynamics. OPA1 and MFN2 were measured as key mitochondrial fusion mediators [50] and Fis1 as a commonly used fission-associated outer-membrane factor [51] that recruits Drp1, the latter involved in constriction and severing of mitochondria [52]. H_2_O_2_ decreased OPA1 and Mfn2 and increased fission-associated proteins (Fig. 6a), including Fis1 and Drp1. NRF1-primed MSC co-culture restored OPA1 and Mfn2 toward CTR levels and attenuated stress-induced fission marker expression (Fig. 6a). To evaluate mitochondrial quality-control pathways, we measured key mitophagy regulators. The PINK1-Parkin axis is a central mitophagy/mitochondrial quality-control pathway crucial for the removal of dysfunctional mitochondria [53, 54]. H_2_O_2_ led to a 5-fold decrease in PINK1 relative to NT. NRF1-primed MSC co-culture led to ∼6-fold change in PINK1 versus SCR and restored Parkin toward baseline (Fig. 6b). Together with PINK1 and Parkin expression, LC3-II, an autophagosome-associated marker used to gauge autophagy/mitophagy engagement [55], was also assessed, with LC3-II levels undergoing a reduction with H_2_O_2_ exposure, and an increase in the NRF1 MSC group (Fig. 6b).

**Figure 6.**
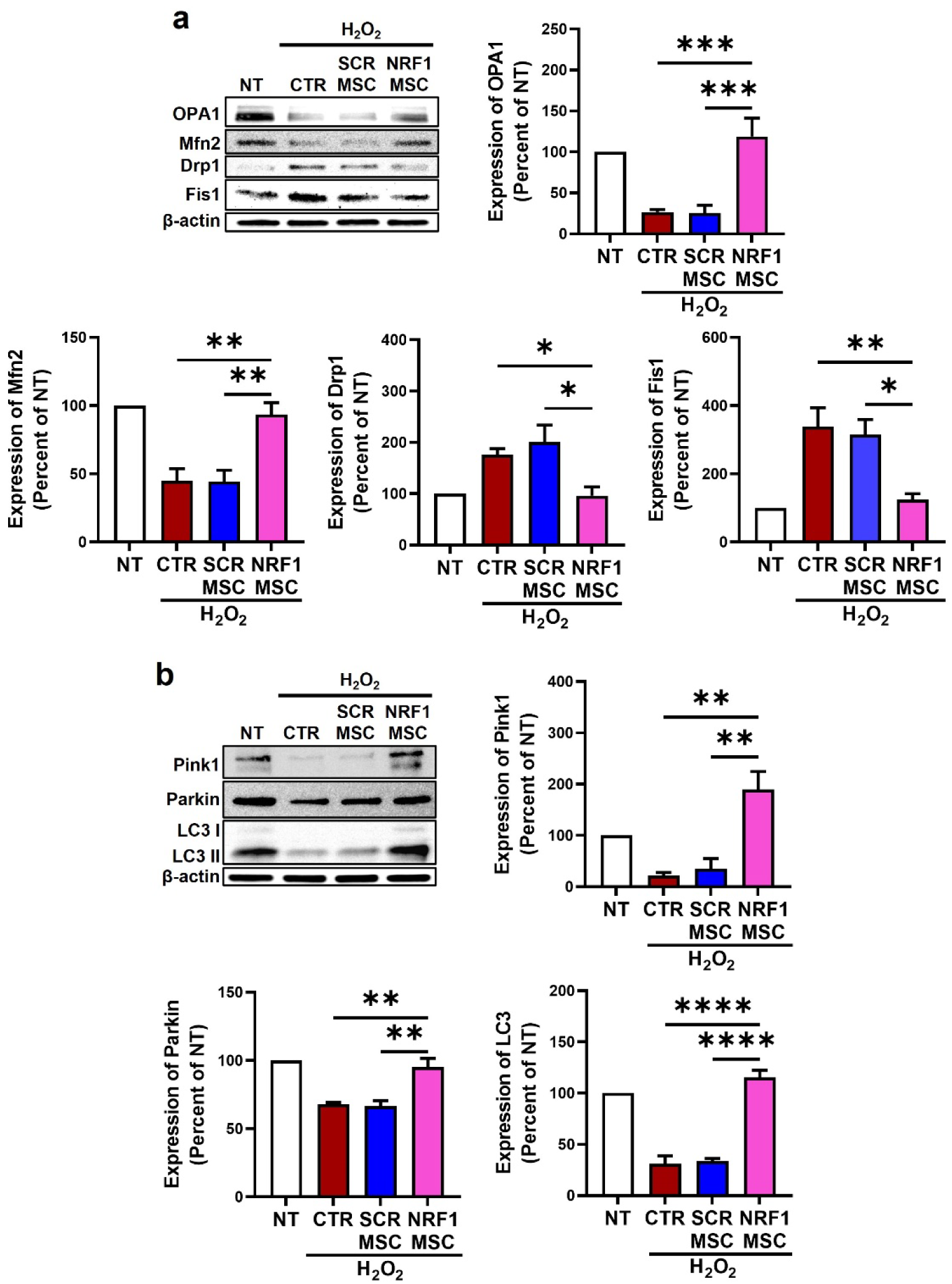
NRF1-primed MSCs restore mitochondrial homeostasis by rebalancing mitochondrial dynamics and regulating mitophagy. H_2_O_2_-exposed (250 µM, 1h) ECs were indirectly co-cultured with MSCs transfected with either scrambled (SCR) or NRF1 mRNA. Controls consisted of healthy, non-treated ECs (NT) and H_2_O_2_-exposed ECs indirectly co-cultured with non-transfected MSCs (CTR). a) Representative western blot and densitometric analysis of expression of proteins involved in fusion (OPA1, Mfn2) and fission (Drp1, Fis1) in ECs. b) Representative western blot and densitometric analysis of expression of mitophagy-related proteins (Pink1, Parkin, LC3 I/II) in ECs. For all Western blots, β-actin was used as a loading control. For densitometric analysis, protein markers were normalized to β-actin expression levels relative to NT. Data are presented as mean ± SEM. *\*p*<0.05, *\*\*p*<0.01, *\*\*\*p*<0.001, *\*\*\*\*p*<0.0001 vs H_2_O_2_ or SCR.

### NRF1-primed MSCs reduce premature senescence and apoptosis in stressed ECs

Oxidative stress and mitochondrial dysfunction contribute to cellular senescence and apoptosis. Staining with SA-β-gal, a marker for senescence, showed that H_2_O_2_ increased the SA-β-gal-positive fraction of cells to ∼45% (Fig. 7a). Co-culture with NRF1-primed MSCs reduced SA-β-gal positivity to ∼18% (Fig. 7a). Subsequently, p53, p21, and p16 were examined because cell-cycle checkpoint/CDK-inhibitor pathways are recognized senescence effectors/biomarkers. Immunoblotting showed that H_2_O_2_ increased p53 (∼4.5-fold change vs NT), p21 (∼3-fold change vs NT), and p16 (∼1.5-fold change vs NT), whereas NRF1-primed MSC co-culture reduced these markers, with p21 and p16 returning toward NT levels (Fig. 7b). Moreover, Annexin V/PI was then selected to distinguish viable vs early apoptotic vs late apoptotic/necrotic cells, while the BAX/Bcl-2 ratio was assessed as a mitochondrial apoptosis rheostat reflecting susceptibility to cell death [56]. Annexin V/PI staining demonstrated increased apoptosis after H_2_O_2_ exposure, which was reduced by NRF1-primed MSC co-culture (Fig. 7c, d). Consistently, H_2_O_2_ increased the BAX/Bcl-2 ratio by ∼2.5-fold change with respect to NT, while NRF1-primed MSC co-culture led to a ∼1.6-fold decrease versus SCR (Fig. 7d).

**Figure 7.**
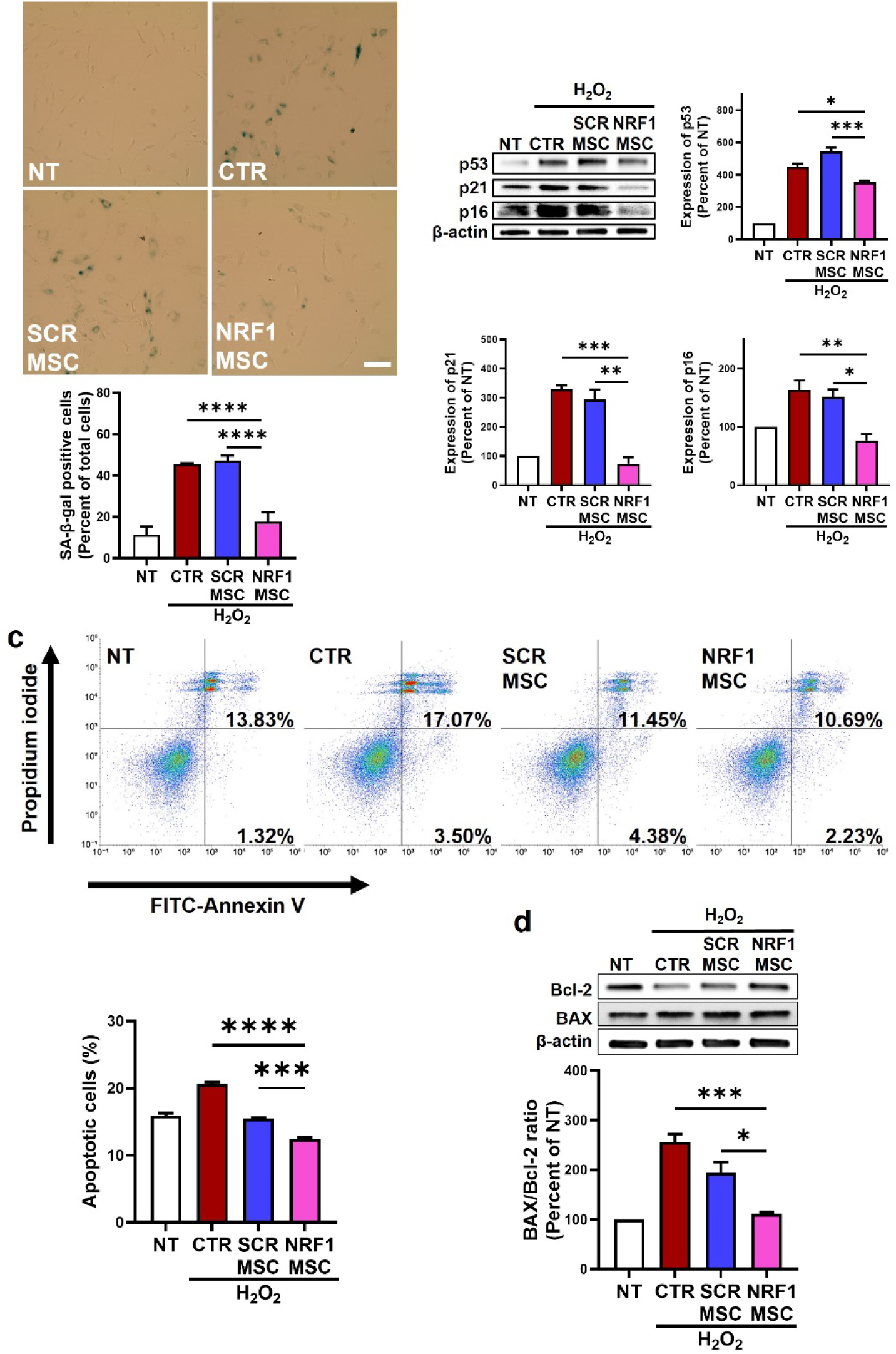
NRF1-primed MSCs prevent oxidative stress-induced premature senescence and apoptosis in ECs. H_2_O_2_-exposed (250 µM, 1 h) ECs were indirectly co-cultured with MSCs transfected with either scrambled (SCR) or NRF1 mRNA. Controls consisted of healthy, non-treated ECs (NT) and H_2_O_2_-exposed ECs indirectly co-cultured with non-transfected MSCs (CTR). a) Representative light microscopy images of SA-β-gal activity in ECs. The bar graph displays SA-β-gal activity, quantified by normalizing the count of cells positive for blue precipitate to the total number of counted cells. Scale bar = 50 µm. b) Representative western blot and densitometric analysis of p53, p21, and p16 expression in ECs. β-actin was used as a loading control. For densitometric analysis, protein markers were normalized to β-actin expression levels relative to NT. c) Representative flow cytometry dot plots displaying percentage of apoptotic cells assessed by Annexin V-FITC and Propidium Iodide (PI) staining. Values in the upper right quadrant (Annexin V^+^/PI^+^) indicate late apoptotic/necrotic cells, while those in the lower right quadrant (Annexin V^+^/PI^-^) represent early apoptotic cells. The bar graph represents the percentage of apoptotic cells, calculated as the sum of early apoptotic and late apoptotic/necrotic populations. d) Representative western blot and densitometric analysis of apoptotic markers (Bcl-2, BAX). β-actin was used as a loading control. For densitometric analysis, protein markers were normalized to β-actin expression levels, and data is expressed as BAX/Bcl-2 ratio. Data are presented as mean ± SEM. \**p*<0.05, \*\**p*<0.01, \*\*\**p*<0.001, \*\*\*\**p*<0.0001 vs H_2_O_2_ or SCR.

### NRF1-primed MSC conditioned medium normalizes stress-associated endothelial dysfunction

To assess functional endothelial outputs, ICAM-1, VCAM-1, and COX2 were measured as established endothelial activation/inflammation-associated markers [57]. H_2_O_2_ increased ICAM-1 (∼3-fold change), VCAM-1 (∼8-fold change), and COX2 (∼5-fold change) compared to NT, whereas NRF1-primed MSCs reduced all three (Fig. 8a). VEGF-A and tube formation assays were then used to examine angiogenic signaling and endothelial morphogenic behavior under stress and rescue conditions. H_2_O_2_ increased VEGF-A (∼1.7-fold change vs NT), whereas NRF1-primed MSCs led to a ∼2-fold decrease in VEGF-A versus SCR (Fig. 8b). In tube formation assays, H_2_O_2_ induced hyper-sprouting with branching points increased (∼15-fold change vs NT). NRF1-primed MSCs significantly reduced branching and stabilized network morphology compared with SCR (Fig. 8c). Lastly, phospho-eNOS (Ser1177) was assessed given that phosphorylation at Ser1177 is a well-supported activation site linked to increased NO output [58, 59], with bradykinin representing a canonical agonist that stimulates endothelial NO signaling. NRF1-primed MSCs restored bradykinin-stimulated eNOS Ser1177 phosphorylation, which was reduced by H_2_O_2_ (Fig. 8c), supporting recovery of NO signaling.

**Figure 8.**
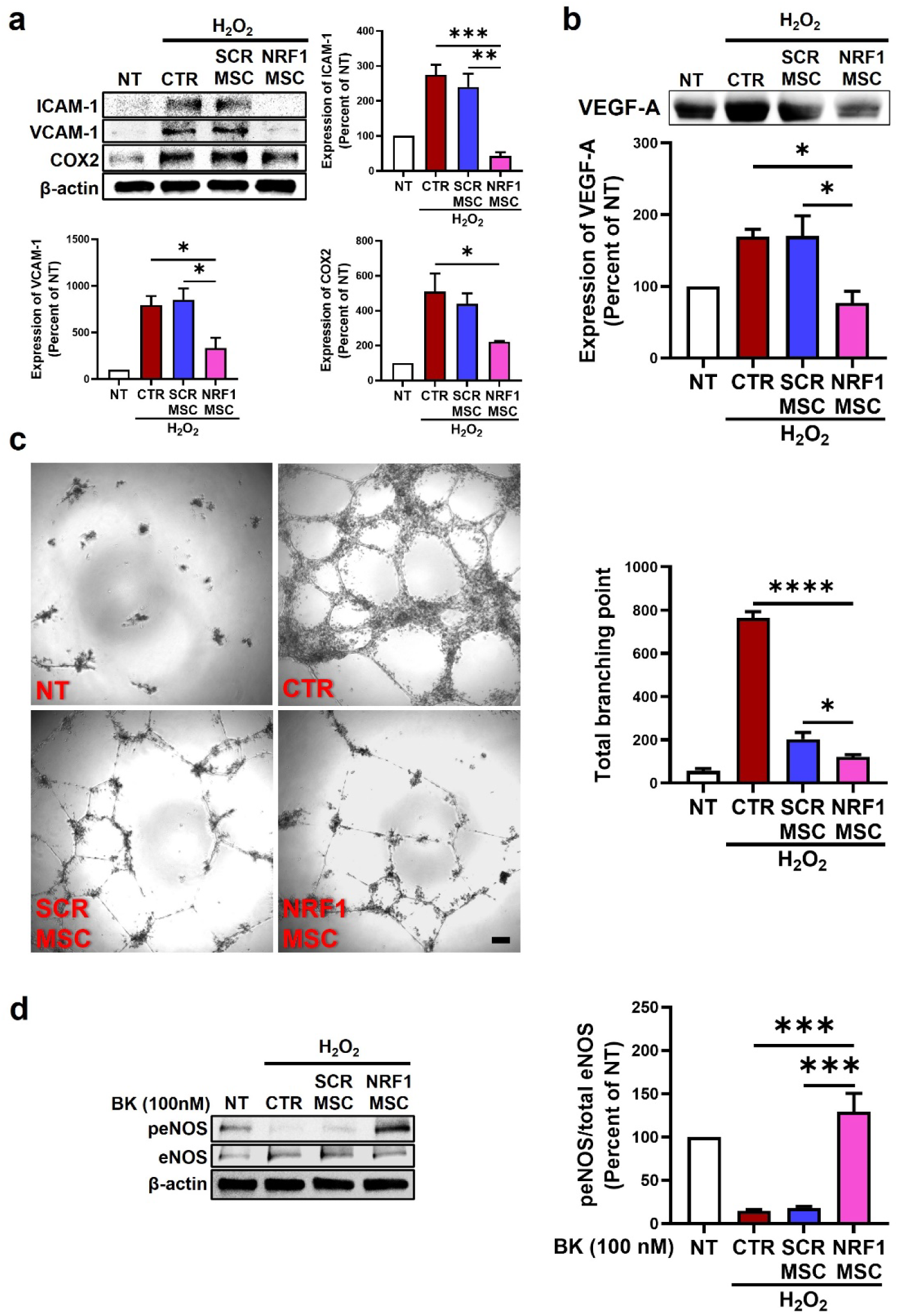
NRF1-primed MSCs ameliorate endothelial dysfunction by suppressing inflammatory angiogenesis and restoring eNOS activity. H_2_O_2_-exposed (250 µM, 1 h) ECs were evaluated using distinct experimental approaches: 1) indirect co-culture with MSCs (Figs. a and d); and 2) treatment with MSC-derived conditioned medium (Fig. c). MSCs were transfected with either SCR or NRF1 mRNA. Control consisted of healthy, non-treated ECs (NT) and H_2_O_2_-exposed ECs (CTR) co-cultured with non-transfected MSCs. a) Representative western blot and densitometric analysis of inflammatory markers (ICAM-1, VCAM-1, COX2) in ECs. b) Representative western blot and densitometric analysis of angiogenic factor VEGF-A present in MSC-derived conditioned medium. c) Representative images of Matrigel tube formation assay and quantification of total branching points. Scale bar = 100 µm. d) Representative western blot and densitometric analysis of phosphorylated and total eNOS in response to 100 nM Bradykinin (BK) stimulation for 30 min. Densitometric analysis was performed to quantify the ratio of phosphorylated eNOS to total eNOS. For all Western blots, β-actin was used as a loading control. For densitometric analysis, protein markers were normalized to β-actin expression levels relative to NT. Data are presented as mean ± SEM. \**p*<0.05, \*\**p*<0.01, \*\*\**p*<0.001, \*\*\*\**p*<0.0001 vs H_2_O_2_ or SCR.

## Discussion

This study establishes NRF1 priming as a coordinated mitochondrial donation program in MSCs that couples enhancement of mitochondrial content with activation of cytoskeletal and vesicular export routes, culminating in functional rescue of oxidative injury in cells. Mitochondrial dysfunction is a convergent driver of endothelial pathology under oxidative stress, where mtROS excess, loss of membrane potential, and impaired OXPHOS destabilize barrier-supporting energetics and amplify inflammatory activation. As we established previously [32], NRF1 overexpression significantly increased mitochondrial biogenesis. Herein, we highlight that NRF1 induction also simultaneously coordinated a mitochondria transfer-ready phenotype, undergoing actin remodeling to favor TNT formation and enhancing secretion of mitochondria-loaded EVs. Functionally, NRF1-primed MSCs increased mitochondrial transfer to oxidatively injured ECs. This increase in mitochondrial delivery aligned with broad restoration of endothelial mitochondrial homeostasis, including improved mitochondrial redox, preserved mitochondrial membrane potential, improved respiration and ATP production, reversal of glycolytic stress reprogramming, rebalanced fusion-fission dynamics, normalization of mitophagy-associated signaling.

Our previous work using transplantation of isolated mitochondria demonstrated that increasing mitochondrial mass can drive substantial functional benefit [7–9]. The current study extends that mitochondria-centric framework by providing a practical alternative to isolating mitochondria from cellular or tissue sources for administration. Specifically, NRF1-primed MSCs act as living mitochondrial depots – mitochondrial hubs capable of transferring mitochondria directly to metabolically compromised cells. This strategy expands the armamentarium of mitochondrial augmentation approaches by leveraging an endogenous, stress-responsive biology (cell-to-cell mitochondrial transfer) while avoiding many limitations associated with mitochondrial isolation, preservation, and delivery.

### NRF1 links mitochondrial biogenesis to a pro-transfer cytoskeletal state and vesicular routing of mitochondria via CD63-positive compartments

NRF1 is classically positioned upstream of mitochondrial biogenesis. Here we show it also couples biogenesis to cytoskeletal remodeling consistent with enhanced intercellular transfer. Upregulation of Cdc42, FAK, and Talin2 aligns with increased actin polymerization, focal-adhesion maturation, and protrusive activity [60], features that can favor formation and stabilization of TNT-like structures. The increase in TNT-like protrusions supports a structural shift toward intercellular connectivity. In parallel, NRF1 also increased Miro1 and EPS8. Miro1 is a key adaptor for mitochondrial transport along cytoskeletal tracks [61], and EPS8 is implicated in actin dynamics and protrusion control [62]. Taken together, our findings support a model in which NRF1 expands the mitochondrial pool while strengthening the infrastructure needed to mobilize and deliver mitochondria via contact-dependent routes.

NRF1 priming also promoted EV-associated routing of mitochondrial material. Increased co-localization of mitochondria with CD63^+^ compartments, enrichment of COXIV in EV isolates, and expansion of AcGFP1^+^/CD63^+^ particles indicate enhanced secretion of mitochondria-containing EVs, notably, without major shifts in vesicle size. Transwell transfer studies further support a contact-independent route consistent with EV-mediated delivery. scRNA-seq independently reinforced this donor-state transition by identifying enrichment of cytoskeletal reorganization, adhesion, and vesicle-trafficking programs in NRF1 MSCs, providing systems-level support for coordinated remodeling rather than a single-effector mechanism. Taken together, these findings position NRF1 as an upstream switch that synchronizes mitochondrial supply with transfer capacity.

Dual-route mitochondrial delivery has direct translational relevance for vascular disease. TNTs enable directed, high-fidelity transfer at sites of cell-to-cell contact, whereas EV-associated delivery supports longer-range, contact-independent transfer – an advantage *in vivo* where donor-recipient proximity varies. NRF1 priming increased transfer to cells in both direct and transwell formats, indicating that enhanced delivery is not confined to a single route, highlighting a redundancy that may prove beneficial.

### NRF1-primed MSCs improve mitochondrial and endothelial function in oxidatively injured ECs

Functionally, increased mitochondrial transfer from NRF1 MSCs translated into measurable improvements in recipient mitochondrial health. NRF1 MSC co-culture reduced mtROS and preserved ΔΨm, consistent with stabilization of respiratory chain integrity and reduced oxidative electron leak. Bioenergetic analyses aligned with rescue of endothelial mitochondrial function. Respiration improved relative to stress conditions, ATP content partially recovered, and stress-associated glycolytic reprogramming. The concurrent downregulation of HIF-1α and glycolytic enzymes (HKII, PFKFB3, LDH-B) suggests that NRF1-dependent mitochondrial support can blunt hypoxia-like signaling induced by oxidative mitochondrial dysfunction. These changes are mechanistically meaningful for vascular biology because endothelial activation and barrier dysfunction are tightly linked to energetic state and redox signaling. The rescue also extended to mitochondrial network organization, with NRF1-primed MSCs rebalancing mitochondrial quality control and mitochondrial dynamics in ECs exposed to oxidative stress. NRF1 MSCs restored fusion mediators (OPA1, MFN2) and attenuated fission-associated markers (Fis1/Drp1), moving away from oxidative stress-induced fragmentation and favoring reformation of an interconnected mitochondrial network needed to distribute mtDNA, respiratory complexes, and metabolites efficiently. In parallel, mitophagy-associated signaling shifted toward normalization, consistent with reduced mitochondrial damage burden and restoration of regulated quality control rather than blunt suppression of turnover pathways. Taken together, enhanced mitochondrial transfer from NRF1 MSCs corresponded to a coherent rescue of endothelial mitochondrial homeostasis. Mitochondrial-level improvements translated into outcomes relevant to vascular pathology, evidenced by reduced senescence and apoptosis, dampened inflammatory activation, normalized stress-associated angiogenic behavior, and recovery of agonist-responsive eNOS Ser1177 phosphorylation, consistent with restoration of endothelial functional state.

### A unified model for translational mitochondrial transfer

To our knowledge, this work is the first to show that direct activation of mitochondrial biogenesis via NRF1 increases intercellular mitochondrial delivery. Prior studies demonstrate that MSCs can transfer mitochondria to endothelium via TNTs to support survival and bioenergetics [11], and emerging work continues to highlight mitochondrial transfer as a driver of tissue repair [63, 64]. Our findings also align with strategies that enhance mitochondrial mass and transfer through stress-responsive pathways (e.g., SIRT1/PGC-1α activation) and emphasize that donor competence depends on cytoskeletal infrastructure [65]. Recent studies further support regulated mitochondria-containing EV secretion via Ca^2+^- and actin-dependent trafficking pathways (e.g., CD38–IP3R–Ca^2+^ signaling) that can boost mitochondria-containing EV output while maintaining donor mitochondrial fitness [36]. Together, these reports reinforce a shared principle: durable benefit is most likely when increased mitochondrial supply is matched to mechanisms that mobilize and export mitochondria.

Our findings data support a two-tier mechanism: 1) donor cell preparation, where NRF1 expands mitochondrial capacity and activates a cytoskeletal/adhesion and trafficking state; and 2) recipient rescue, where delivered mitochondrial material restores mitochondrial function, rebalances dynamics and quality control, improves energy production, and prevents stress-driven senescence and apoptosis, culminating in improved endothelial functional outputs. Our novel hub concept is translationally attractive for several reasons. First, NRF1 mRNA priming is transient and avoids stable genetic modification, making it compatible with manufacturing workflows as a short preconditioning step before cell administration or EV harvesting. Second, because NRF1 priming enhances mitochondrial EV output, the approach is compatible with cell-based and cell-free therapeutic strategies, including EV-exclusive therapies aimed at mitochondrial support. Third, our data reinforce the value of targeting mitochondrial networks and upstream transcriptional regulators (e.g., NRF1-driven biogenesis, fusion-fission balance, bioenergetics) rather than attempting to modulate individual downstream stress pathways in isolation. Vascular disease states typically involve simultaneous disruption of redox balance, respiration, mitochondrial dynamics, and quality control. A systems-level strategy that rebuilds mitochondrial competence and restores transfer capacity may therefore be better aligned with the complexity of oxidative-stress vasculopathies.

## Conclusions

NRF1 priming converts MSCs into mitochondrial donor hubs for stressed endothelium by increasing mitochondrial supply and activating the machinery required for intercellular delivery. NRF1 synchronizes mitochondrial biogenesis with cytoskeletal remodeling and vesicular export, enabling both contact-dependent and contact-independent mitochondrial transfer. This donor-state transition translates into restoration of endothelial mitochondrial integrity, metabolic balance, and function after oxidative injury.

## Acknowledgements

The authors are grateful for funding from the Houston Methodist Research Institute. The authors also thank David L. Haviland and Elizabeth Jardinella from the Flow Cytometry Core at Houston Methodist Research Institute for technical support and assistance. The authors thank Bruna Corradetti (Baylor College of Medicine [BCM]) for use of ZetaView instrument. The authors acknowledge the Mouse Metabolism and Phenotyping Core (MMPC) at BCM. All schematics were created with BioRender.com.

