## Supplementary Materials for "Enhanced intercellular transfer of mitochondria from nuclear respiratory factor 1 (NRF1)-primed mesenchymal stem cells: towards creation of superior mitochondrial delivery hubs"

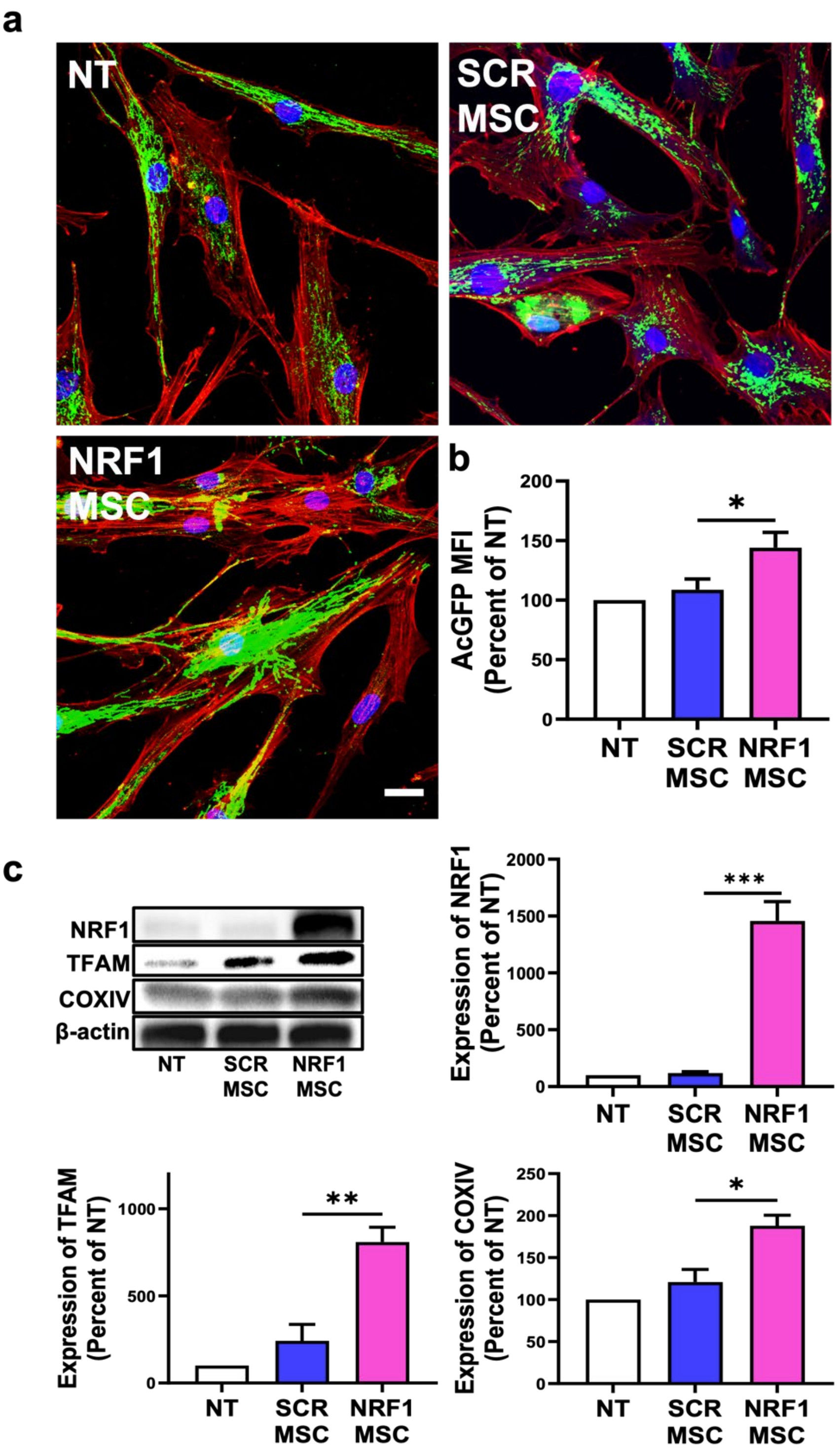


**Figure S1. NRF1 overexpression promotes mitochondria biogenesis and upregulates downstream protein expression.** MSCs were transfected with either scrambled (SCR) or NRF1 mRNA. Controls consisted of non-transfected MSCs (NT). a) Representative confocal images of MSCs. Mitochondria are visualized by AcGFP1 (green), F-actin (red), and DAPI (Nuclei). Scale bar = 25 µm. b) Quantification of AcGFP1 mean fluorescence intensity (MFI) by flow cytometry. c) Representative western blot of NRF1, TFAM, and COXIV protein expression. β-actin was used as a loading control. Densitometric analysis was performed to quantify protein expression. Protein markers were normalized to β-actin expression levels relative to NT. **p*<0.05, ***p*<0.01, ****p*<0.001 vs SCR.

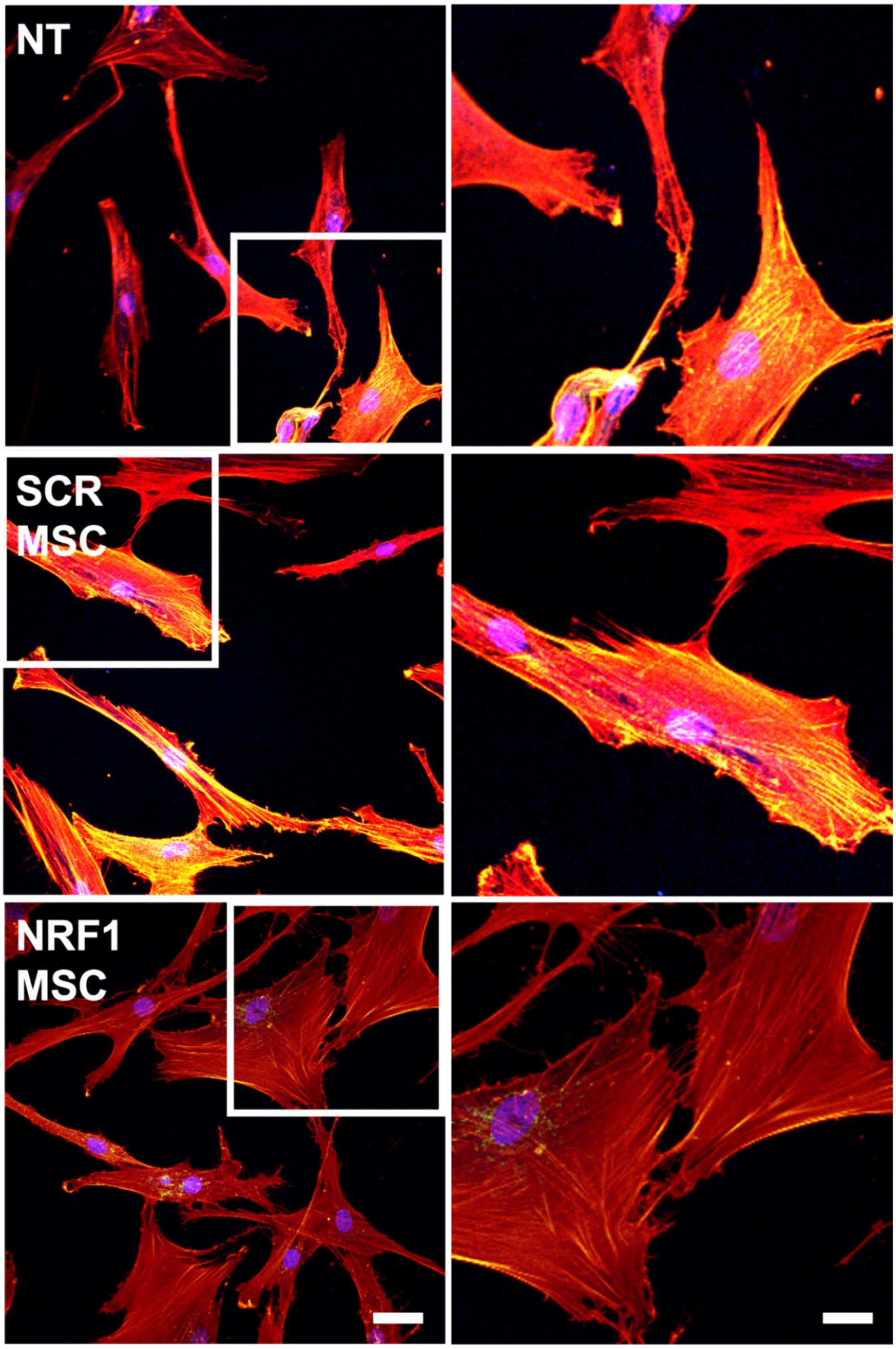


**Figure S2. Representative single optical sections from z-stack confocal imaging of MSC tunneling nanotube (TNT) formation.** MSCs were transfected with either scrambled (SCR) or NRF1 mRNA. Controls consisted of non-transfected MSCs (NT). Representative confocal microscopy images of MSCs stained for F-actin (red) and nuclei (blue). Scale bar = 12.5 µm. These images are representative single optical sections selected from z-stack layered images used to verify and to count TNT number. Boxed regions indicate magnified regions (2X magnification) and are images presented in Figure 1c as monochromatic for clarity of presentation. Scale bar = 25 µm.

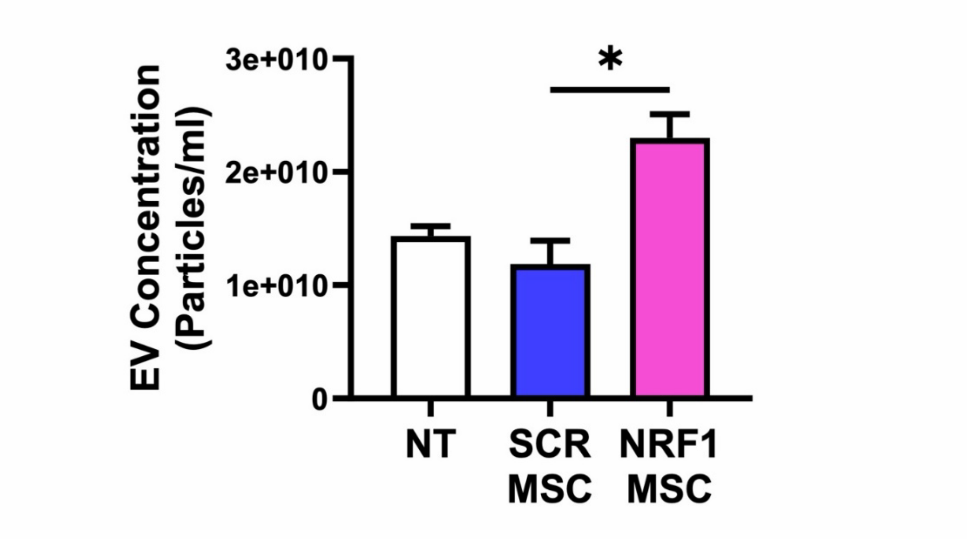


| EV Concentration (Particles/ml) | NT | SCR MSC | NRF1 MSC |
|---|---|---|---|
| | 1.43E+010 ± 1.53E+009 | 1.19E+010 ± 3.59E+009 | 2.30E+010 ± 3.61E+009 |

**Figure S3. Quantification of EV concentration.** MSCs were transfected with either scrambled (SCR) or NRF1 mRNA. Controls consisted of non-transfected MSCs (NT). MSC-derived EV concentration was determined using the Zetaview system. *$p$<0.05 vs SCR

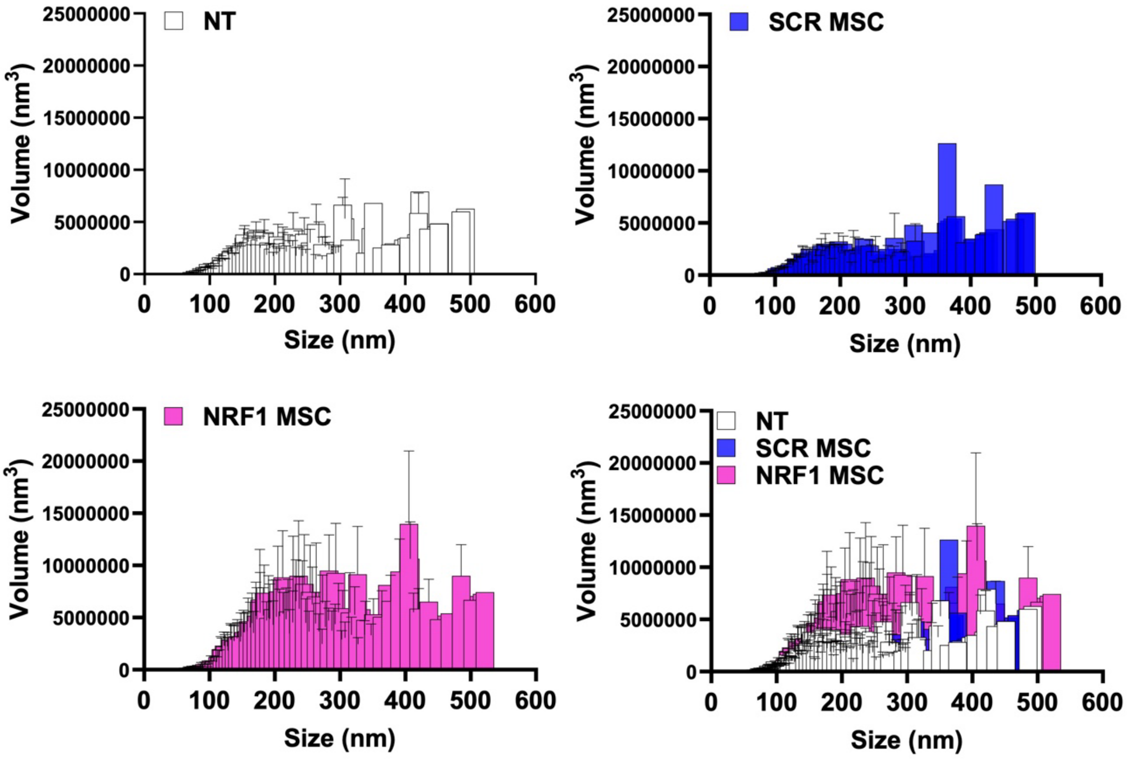


**Figure S4. Characterization of EV size distribution.** MSCs were transfected with either scrambled (SCR) or NRF1 mRNA. Controls consisted of non-transfected MSCs (NT). Representative individual histograms and overlay of the volumetric distribution of MSC-derived EV size.

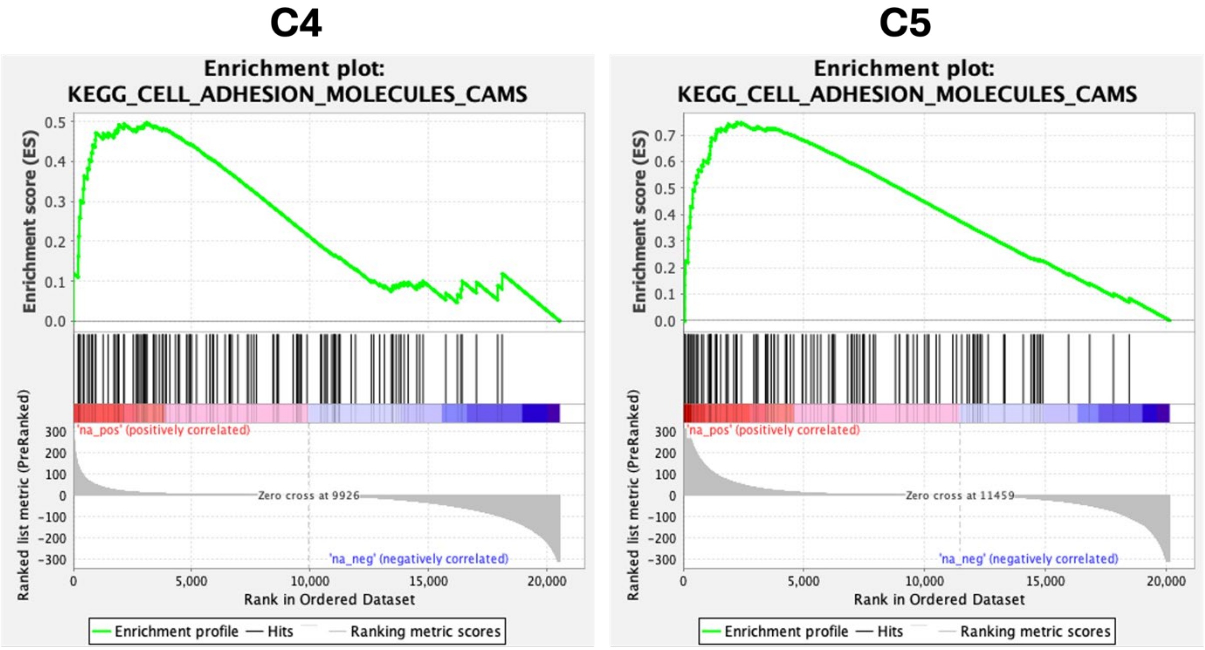


**Figure S5. GSEA enrichment plots of the most significant adhesion molecule pathway in NRF1-transfected MSC dominant subpopulations.** GSEA was performed on Clusters 4 and 5, which mainly comprise NRF1-transfected MSCs. Plots display the KEGG_CELL_ADHESION_MOLECULES_CAMS gene set, which was identified as the top-ranked pathway within the adhesion molecule related pathway.

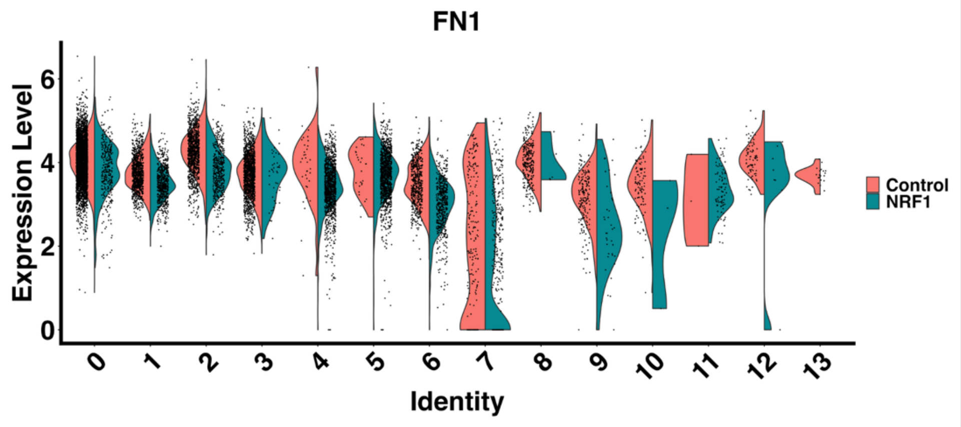


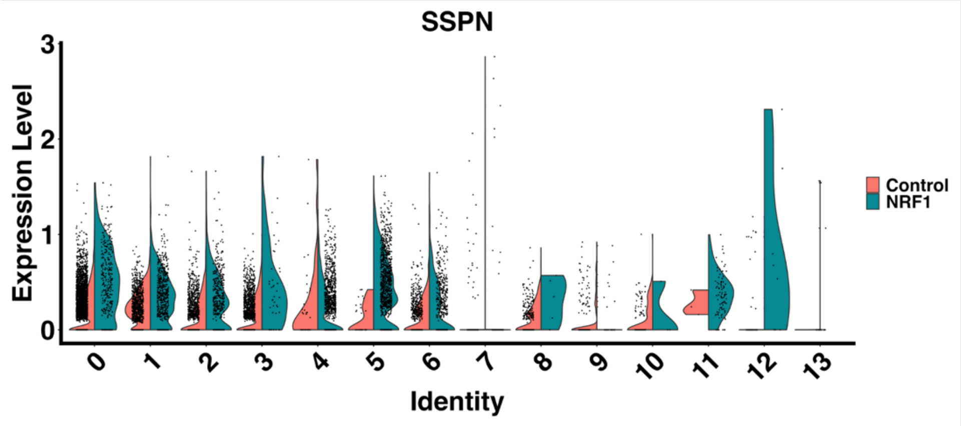


**Figure S6. Expression profiles of the top two significantly upregulated cell adhesion genes in NRF1-transfected MSC dominant subpopulations.** Violin plots display the expression levels of FN1 and SSPN, genes identified as the two most significantly upregulated markers among cell-cell adhesion related genes in Clusters 4 and 5, which represent the predominant subpopulations in the NRF1-transfected group. Red: CTR and SCR-transfected MSCs group; Blue: NRF1-transfected MSCs.

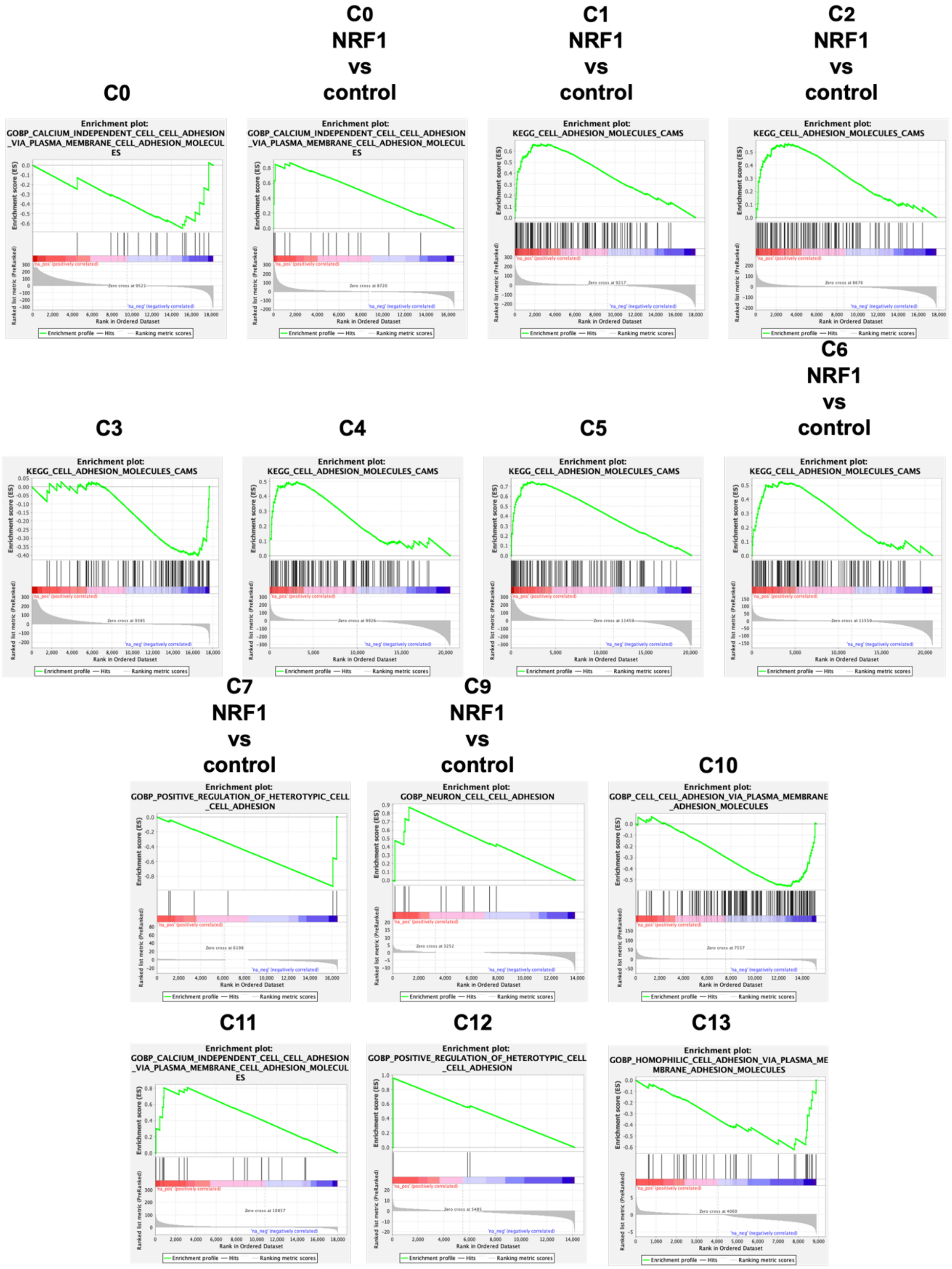


**Figure S7. Landscape of the most significant cell adhesion pathways enriched across different MSC subpopulations.** GSEA enrichment plots display the top-ranked cell adhesion pathways for each individual cluster. The KEGG_CELL_ADHESION_MOLECULES_CAMS pathway is consistently enriched across multiple clusters (C1; NRF1vsCTR, C2; NRF1vsCTR, C3, C6; NRF1vsCTR). CALCIUM_INDEPENDENT_CELL_CELL_ADHESION (C0, C0; NRF1vsCTR, C11), GOBP_POSITIVE_REGULATION_OF_HETEROTYPIC_CELL_CELL_ADHESION (C7; NRF1vsCTR, C12), GOBP_NEURON_CELL_CELL_ADHESION (C9; NRF1vsCTR), and GOBP_HOMOPHILIC_CELL_ADHESION_VIA_PLASMA_MEMBRANE_MOLECULES (C13).

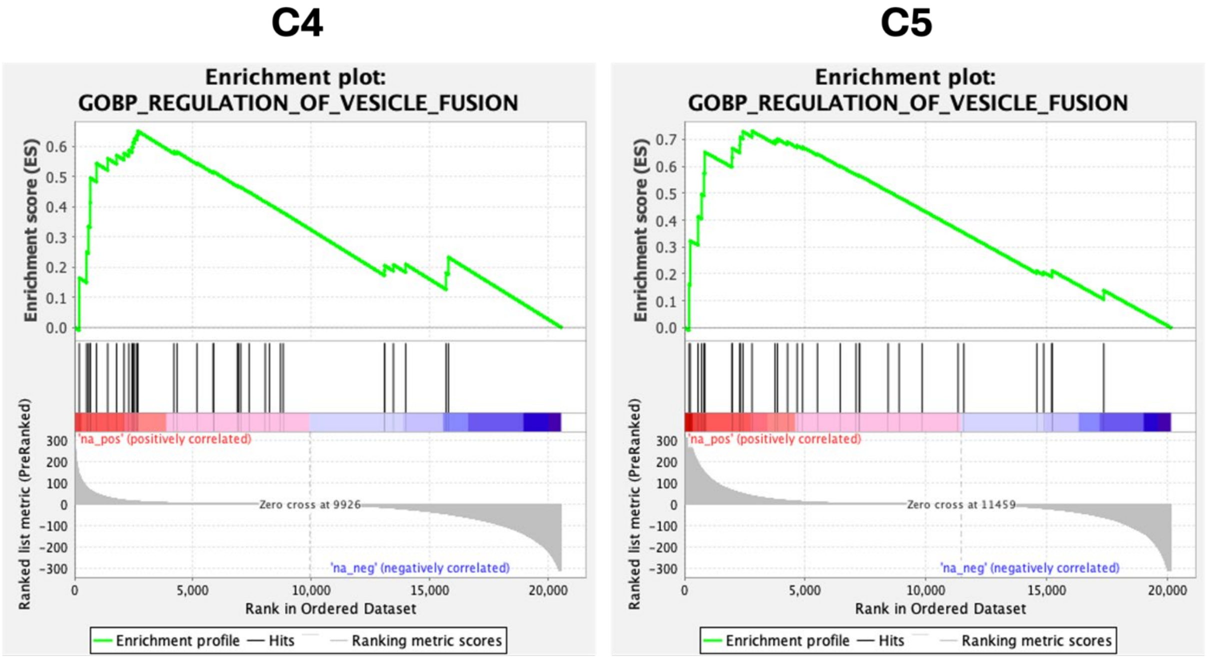


**Figure S8. GSEA enrichment plots of the most significant EV-related pathway in NRF1-transfected MSC dominant subpopulations.** Gene Set Enrichment Analysis (GSEA) was performed on Clusters 4 and 5, which mainly comprise NRF1-transfected MSCs. Plots display the GOBP_REGULATION_OF_VESICLE_FUSION gene set, which was identified as the top-ranked pathway within the EV related pathway.

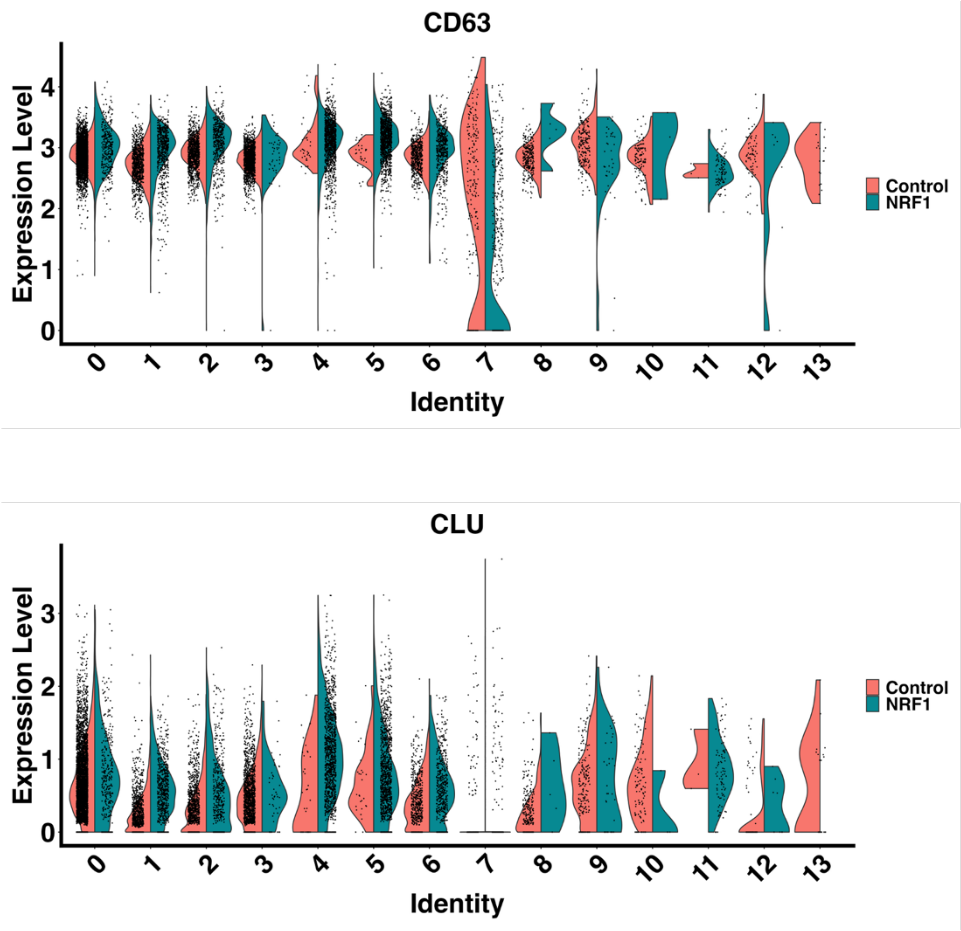


**Figure S9. Expression profiles of the top two significantly upregulated EV-related genes in NRF1-transfected MSC dominant subpopulations.** Violin plots display the expression levels of CD63 and CLU, genes identified as the two most significantly upregulated markers among EV related genes in Clusters 4 and 5, which represent the predominant subpopulations in the NRF1-transfected group. Red: CTR and SCR-transfected MSCs; Blue: NRF1 transfected MSCs.

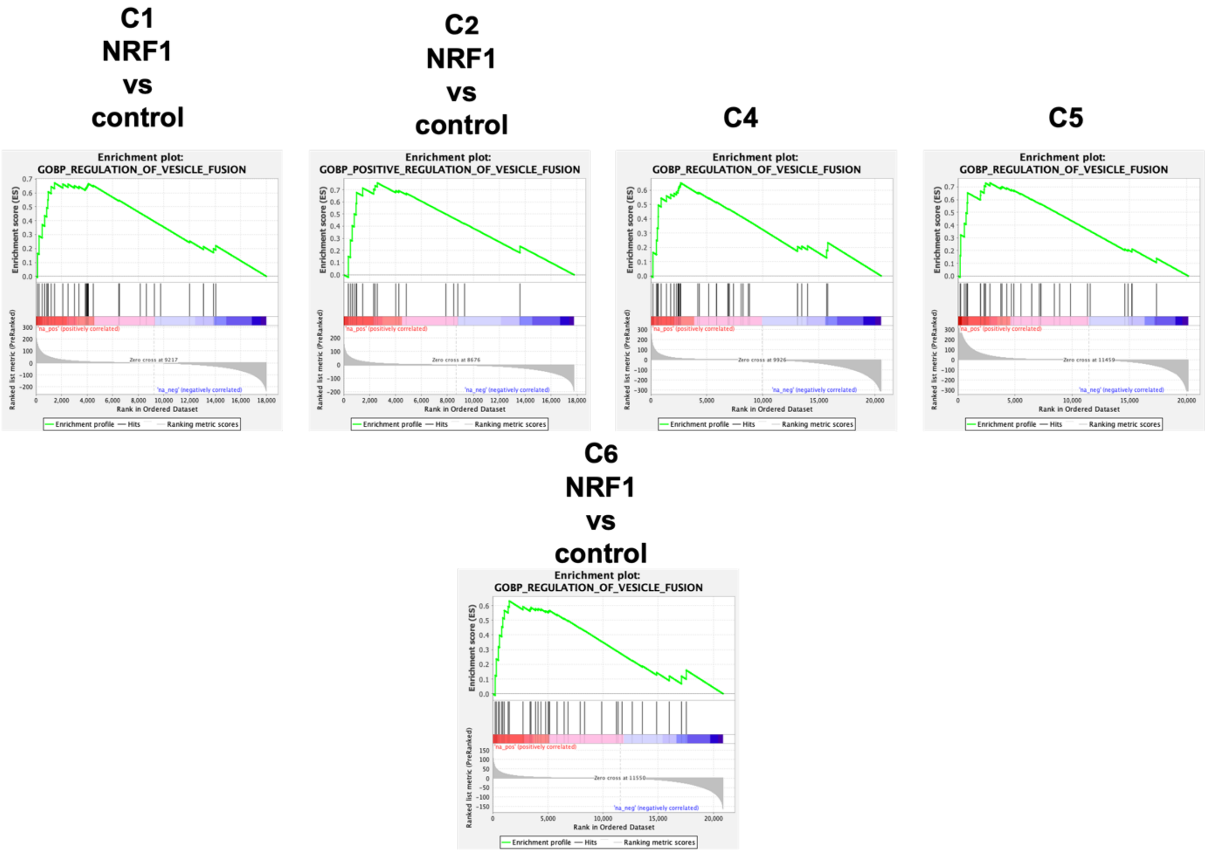


**Figure S10. Landscape of the most significant vesicle fusion pathways enriched across different MSC subpopulations.** GSEA enrichment plots display the top-ranked EV-related pathways for each individual cluster. The GOBP_REGULATION_OF_VESICLE_FUSION pathway is consistently enriched across multiple clusters (C1; NRF1vsCTR, C4, C5, C6; NRF1vsCTR). GOBP_POSITIVE_REGULATION_OF_VESICLE_FUSION (C2; NRF1vsCTR).

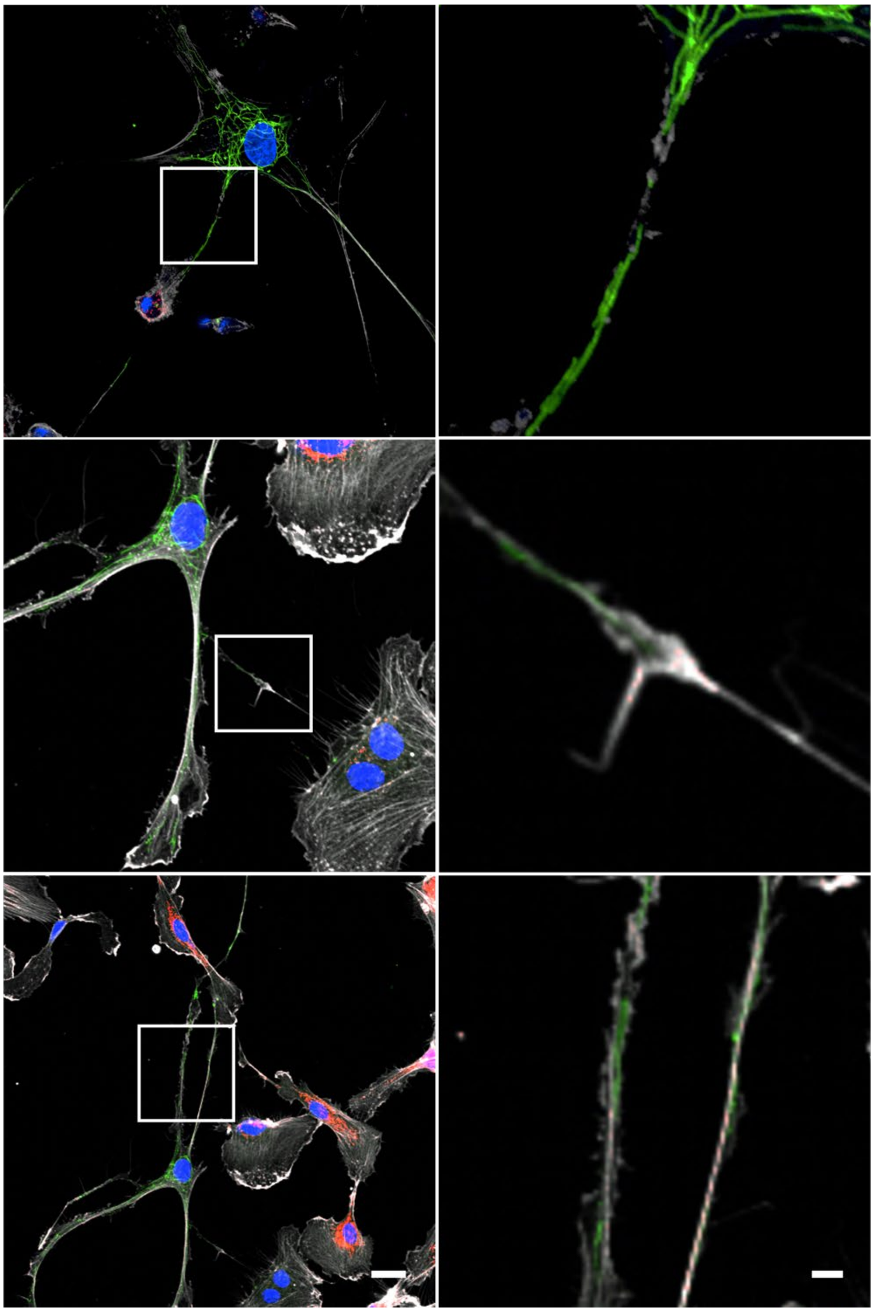

**Figure S11. Visualization of TNT-mediated mitochondrial donation from NRF1-primed MSCs to ECs.** $H_2O_2$-exposed (250 µM, 1 h) ECs were co-cultured directly with MSCs transfected with either scrambled (SCR) or NRF1 mRNA. Representative confocal microscopy images demonstrating direct transfer of mitochondria. MSC mitochondria-AcGFP1 (green), F-actin (gray), EC mitochondria-mCherry (red), and DAPI (Nuclei). Scale bar = 25 µm. Boxed regions represent magnified regions to the left of the images. Scale bar = 5 µm.

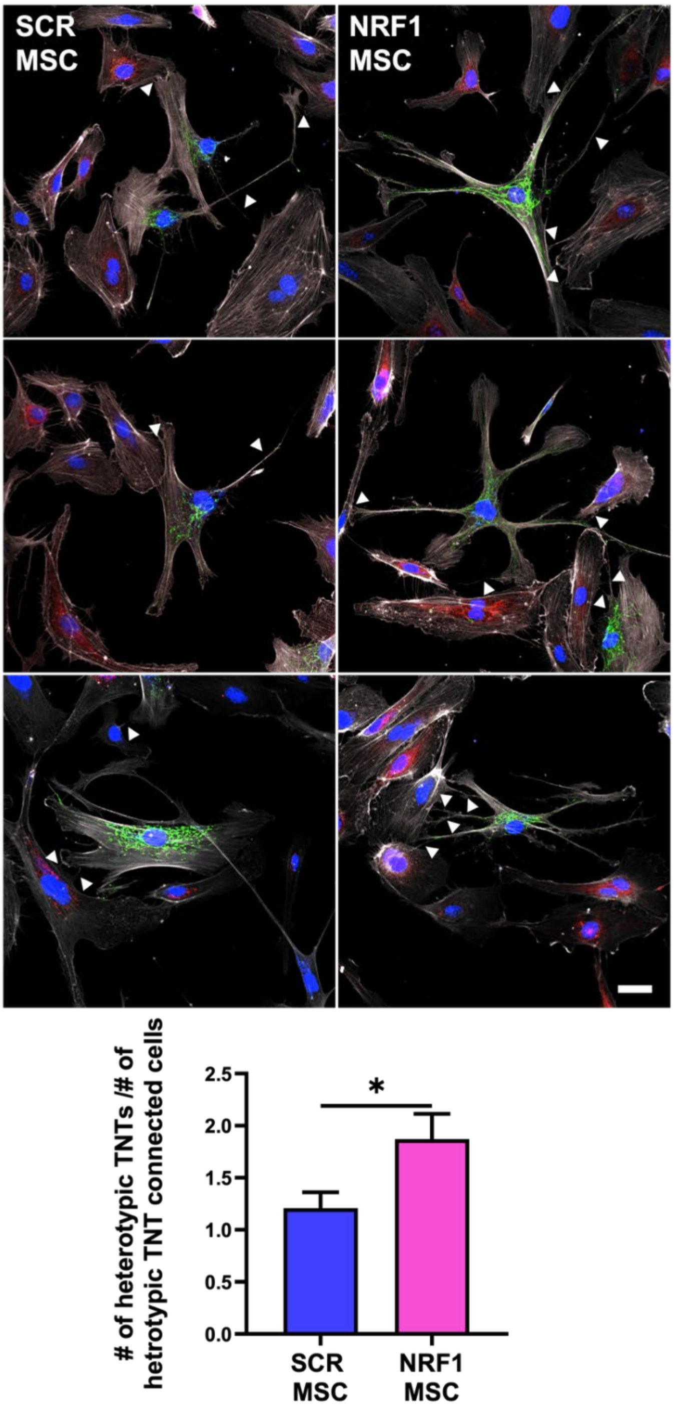


**Figure S12. NRF1 overexpression promotes TNT formation between MSCs and ECs exposed to oxidative stress.** $H_2O_2$-exposed (250 μM, 1 h) ECs were co-cultured directly with MSCs transfected with either scrambled (SCR) or NRF1 mRNA. Representative images showing intercellular connections. White arrowheads indicate TNT formation between MSCs and ECs. MSC mitochondria-AcGFP1 (green), F-actin (gray), EC mitochondria-mCherry (red), and DAPI (Nuclei). Scale bar = 25 μm. The bar graph represents the number of heterotypic TNTs per the number of heterotypic TNT connected MSCs and ECs. **p*<0.05 vs SCR.

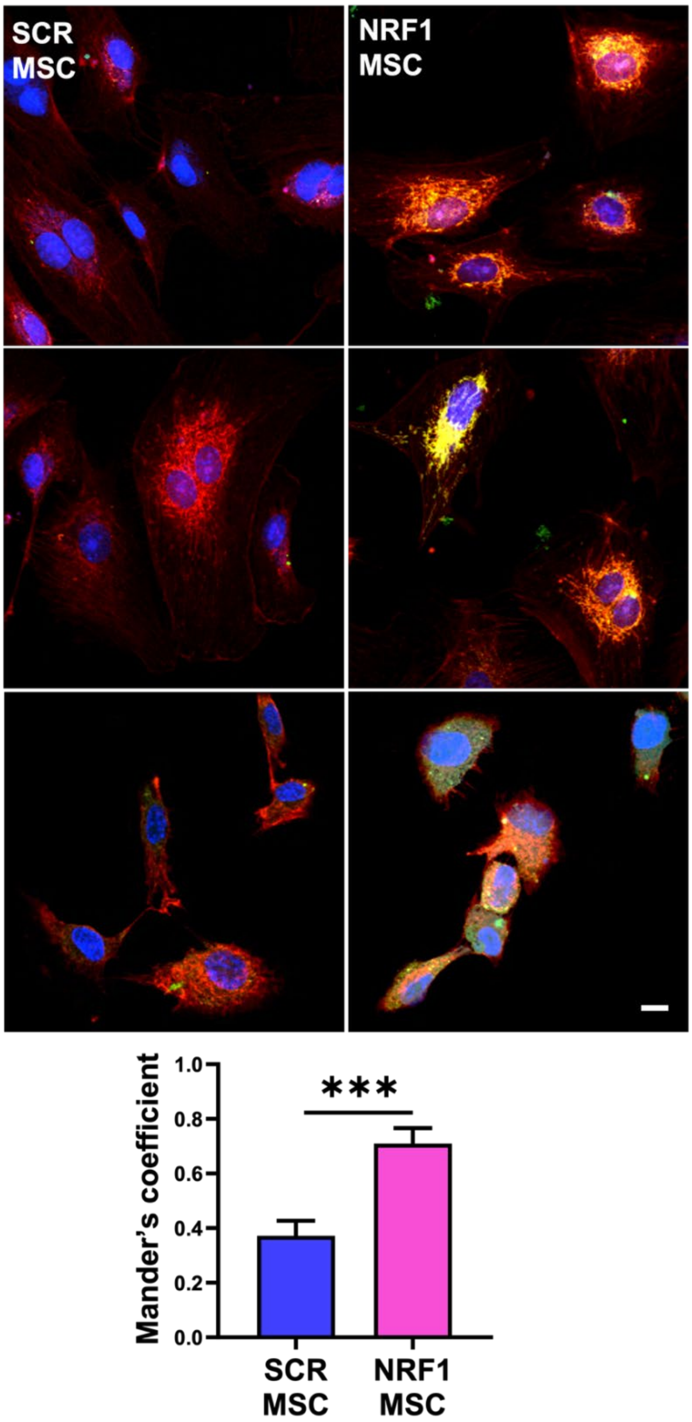


**Figure S13. Colocalization of EV-delivered, MSC-derived mitochondria with mitochondria in recipient ECs.** $H_2O_2$-exposed (250 µM, 1 h) ECs were co-cultured indirectly with MSCs transfected with either scrambled (SCR) or NRF1 mRNA. Representative confocal microscopy images of mitochondrial colocalization in ECs. MSC mitochondria-AcGFP1 (green), F-actin (gray), EC mitochondria-mCherry (red), and DAPI (Nuclei). Yellow indicates colocalized mitochondria. Scale bar=25 µm. The bar graph indicates quantitative analysis of mitochondrial colocalization using Mander's overlap coefficient. ****p*<0.001 vs SCR
